# Plasma membrane PI4P recruits SNAP47 to mediate SNARE-dependent AMPAR exocytosis during synaptic potentiation

**DOI:** 10.64898/2026.08.08.743640

**Authors:** Jing Xi, Shen Wang, Deng Pan, Yanrui Yang, Ranran Mao, Sin Man Lam, Wenting He, Guanghou Shui, Yang Niu, Lei Chen, Cong Ma, Jia-Jia Liu

## Abstract

The synaptic delivery of the AMPA-type glutamate receptors (AMPARs) is crucial for longterm potentiation (LTP) of excitatory synapses, yet the mechanisms underlying neuronal activity-dependent AMPAR exocytosis at the plasma membrane (PM) remain unclear. We previously demonstrated that the PM synthesis of the phosphoinositide PI4P is enhanced upon LTP induction and that PM PI4P, not PI(4,5)P_2_, is required for activity-induced AMPAR exocytic trafficking. Here, we show that AMPARs are exocytosed at PI4P-enriched dendritic PM microdomains in potentiated hippocampal neurons. The Q-SNARE SNAP47 binds PI4P via its pleckstrin homology (PH)-like domain. This interaction recruits SNAP47 to the PM, promoting the exocytic fusion of AMPAR vesicles through the SNAP47–Syntaxin-3–VAMP2 SNARE complex. In the hippocampus, the SNAP47–PI4P interaction is necessary for both LTP and long-term memory. Our findings reveal a mechanistic role for PI4P in mediating activity-dependent, SNARE-driven fusion of AMPAR exocytic vesicles with the PM.

## Introduction

In the brain, α-amino-3-hydroxy-5-methyl-4-isoxazolepropionic acid-type ionotropic glutamate receptors (AMPARs) mediate the majority of fast excitatory synaptic transmission. Changes in AMPAR numbers at postsynaptic sites represent a key component of synaptic plasticity, the cellular correlate of learning and memory ^1^. During N-methyl-D-aspartate receptor (NMDAR)-mediated LTP, Ca^2+^ signaling stimulates exocytic trafficking of AMPAR transport vesicles primarily to extrasynaptic sites on the PM of dendritic shafts adjacent to potentiated spines ^2,3^. Subsequently, receptor molecules reach the postsynaptic membrane at the spine head via lateral diffusion and are stabilized by scaffold and auxiliary proteins ^4^, leading to sustained increase in synaptic strength. Although mechanisms underlying AMPAR synaptic trafficking have been extensively studied, the molecular basis of activity-dependent AMPAR exocytosis remains poorly understood.

Electrophysiological studies of hippocampal CA1 neurons have revealed that Ca^2+^-triggered membrane fusion events in postsynaptic cells contribute to LTP ^3,5^. The soluble *N*-ethylmaleimide-sensitive factor attachment protein receptor (SNARE) family of proteins mediates membrane fusion between organelles and the PM ^6^. The Ca^2+^ sensors synaptotagmin-1 (Syt-1) and synaptotagmin-7 (Syt-7), together with their co-factor complexin, regulate SNARE-mediated membrane fusion and are essential for activity-dependent AMPAR exocytosis ^3,7^. Several SNARE proteins—the R-SNARE vesicle-associated membrane protein 2 (VAMP2, also known as synaptobrevin 2, a vesicular [v]-SNARE), the Q-SNAREs synaptosomal-associated protein 47 (SNAP47) and the PM-localized syntaxin-3 (Stx3, a target [t]-SNARE)—are implicated in regulated exocytosis of AMPARs during LTP ^8–10^. However, how neuronal activity regulates SNARE complex recruitment and assembly to mediate fusion of AMPAR transport vesicles with the PM remains unclear.

Phosphoinositides (or phosphatidylinositol phosphates, PIPs) are low-abundance phospholipids generated by phosphorylation of phosphatidylinositol (PI) at the 3, 4, and 5 positions of the inositol ring. They are differentially distributed in the cytoplasmic leaflet of the PM and organelle membranes of eukaryotic cells, where they regulate diverse cellular processes by recruiting effector proteins involved in membrane remodeling, fusion and fission, vesicle trafficking, and non-vesicular lipid transport at membrane contact sites ^11,12^. Notably, phosphoinositide metabolism is dysregulated in aging and diseased brains ^13–15^, linking these lipids to synaptic function. Among the seven phosphoinositides, phosphatidylinositol 4-phosphate (PI4P) and phosphatidylinositol 4,5-bisphosphate (PI(4,5)P_2_) are the most abundant and are relatively enriched in the PM. Previous studies have shown that phosphatidylinositol 3,4,5-trisphosphate (PI(3,4,5)P_3_) and PI(4,5)P_2_ function in synaptic vesicle trafficking ^16–18^. Most recently, it was reported that PM PI4P facilitates synaptic vesicle exo-endocytosis to sustain efficient synaptic transmission ^19^. Although the role of PI(4,5)P_2_ in vesicle exocytosis and SNARE-mediated membrane fusion has been well established ^20–24^, the precise molecular function(s) of PI4P in exocytic trafficking of synaptic cargoes is largely unknown.

We previously reported that LTP stimuli induce rapid increases in PM PI4P levels in rodent hippocampal neurons, and that PM PI4P is required for activity-dependent exocytic trafficking of AMPAR vesicles ^25^. In activated neurons, in response to elevated cytosolic Ca^2+^, the membrane tethering protein extended synaptotagmin 1 (E-Syt1) recruits PI4KIIIα to endoplasmic reticulum (ER)‒PM contact sites to catalyze PM synthesis of PI4P ^26^. Intriguingly, depletion of PM PI(4,5)P_2_ has no impact on LTP stimulus-induced AMPAR surface expression ^25^, raising the question of whether PI4P plays a direct role in SNARE-mediated fusion of AMPAR vesicles with the PM. In this study, we investigated the molecular mechanism for PI4P-mediated AMPAR exocytosis. We show that PI4P recruits SNAP47 via its PH-like domain to the PM to facilitate exocytic fusion of AMPAR vesicles mediated by the SNAP47–Stx3–VAMP2 SNARE complex. We further demonstrate that the SNAP47–PI4P interaction is required for LTP expression and long-term memory *in vivo*.

## Results

### SNAP47 and Stx3 mediate activity-dependent AMPAR exocytosis and complex with VAMP2

Although the role of PM PI(4,5)P_2_ in exocytosis has been well established, previously we found that inhibition of PI4KIIIα-catalyzed PM PI4P synthesis, rather than acute depletion of PM PI(4,5)P_2_ by G protein-coupled receptor (GPCR)-activated, phospholipase C-catalyzed hydrolysis, abolishes activity-induced AMPAR exocytic trafficking in neurons ^25^. As dendritic PM levels of PI4P are rapidly upregulated upon LTP induction ^25^, we hypothesized that PI4P could play a direct role in regulating SNARE-mediated fusion of AMPAR vesicles. To test this possibility, we first examined the spatiotemporal relationship between changes in PM PI4P and AMPAR exocytosis in potentiated hippocampal neurons by total internal reflection fluorescence (TIRF) microscopy-based live imaging. Fusion of AMPAR transport vesicles with the PM is detected by dequenching of pHluorin, a pH-sensitive GFP variant fused to the extracellular domain of the AMPAR subunit GluA1 (SEP-GluA1) ^27^. Shortly after application of the NMDAR co-agonist glycine, we observed repeated exocytosis of SEP-GluA1 at regions of the dendritic PM exhibiting increased fluorescence of the PI4P biosensor mCherry-PH^OSBP^. In contrast, SEP-GluA1 exocytic signals did not overlap with those of mCherry-PH^PLCδ1^, the PI(4,5)P_2_ probe (Fig. 1A and Video 1). These findings suggest that PI4P-enriched PM microdomains serve as hotspots for AMPAR exocytosis.

**Fig. 1.**
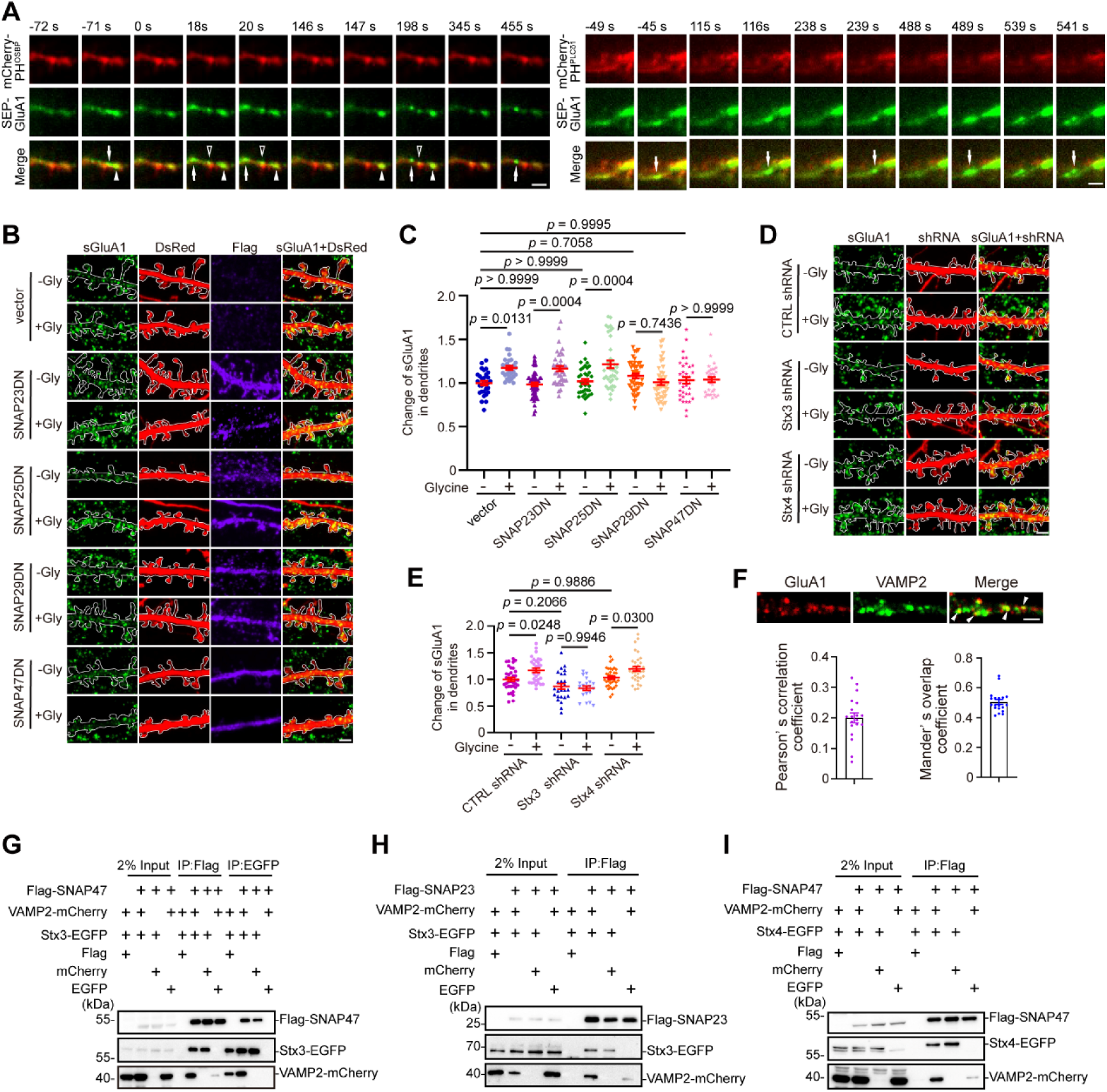
SNAP47 and syntaxin-3 are required for activity-dependent AMPAR exocytosis and complex with VAMP2. (A) Cultured rat hippocampal neurons were co-transfected on DIV12 with constructs expressing SEP-GluA1 and mCherry-PH^OSBP^ or mCherry-PH^PLCδ1^. On DIV16, chemical LTP (cLTP) was induced by glycine application, and neurons were imaged live using TIRF microscopy. Shown are representative images of dendrites with AMPAR exocytosis. Arrows and arrowheads indicate SEP-GluA1 exocytic events at the plasma membrane. The arrowheads indicate co-localization of the mCherry-PH^OSBP^ signal with hotspots of SEP-GluA1 exocytosis. Scale bars, 2 μm. (B) Cultured mouse hippocampal neurons were co-transfected on DIV11 with constructs expressing DsRed and a Flag-tagged dominant-negative (DN) mutant of SNAP23, SNAP25, SNAP29 or SNAP47. On DIV16, cLTP was induced by glycine application for 5 min. Neurons were fixed 5 min following glycine washout and immunostained for Flag, RFP, and surface-expressed GluA1 (sGluA1). Shown are representative confocal fluorescence images. Scale bar, 2 μm. (C) Quantification of surface GluA1 in dendrites in (B). Data represent mean ± SEM, n = 29-47 neurons from two independent experiments, 1-3 dendrites/neuron. Statistical significance was determined by one-way ANOVA analysis of variance with a Tukey post hoc test. (D) Cultured mouse hippocampal neurons were transfected on DIV11 with constructs co-expressing GFP and non-targeting control shRNA (CTRL shRNA) or shRNA targeting Stx3 (Stx3 shRNA) or Stx4 (Stx4 shRNA). On DIV16, cLTP was induced by glycine application for 5 min. Neurons were fixed 5 min following glycine washout and immunostained for GFP and surface-expressed GluA1. Shown are representative confocal microscopy images. Scale bar, 2 μm. (E) Quantification of surface GluA1 in dendrites in (D). Data represent mean ± SEM, n = 21-37 neurons from two independent experiments, 1-3 dendrites/neuron. Statistical significance was determined by one-way ANOVA analysis of variance with a Tukey post hoc test. Scale bars, 2 μm. (F) DIV17 mouse hippocampal neurons were immunostained for intracellular GluA1 and VAMP2. Shown are representative confocal images (arrowheads indicate colocalized signals) and quantification of colocalization between GluA1 and VAMP2 in dendrites. Data represent mean ± SEM, n = 19 neurons, 1-3 dendrites/neuron. Scale bar, 2 μm. (G) HEK293T cells co-transfected with constructs expressing Flag-SNAP47, VAMP2-mCherry, and Stx3-EGFP were lysed and subjected to immunoprecipitation (IP) using anti-Flag or anti-GFP antibody-conjugated agarose beads. Input and bound fractions were analyzed by SDS‒PAGE followed by immunoblotting with antibodies against Flag, GFP, and RFP. The Flag vector served as control for Flag-SNAP47, mCherry served as control for VAMP2-mCherry, and EGFP served as control for Stx3-EGFP. (H) HEK293T cells co-transfected with constructs expressing Flag-SNAP23, VAMP2-mCherry, and Stx3-EGFP were lysed and subjected to IP with anti-Flag beads. (I) Same as (H) except that cells were co-transfected with constructs expressing Flag-SNAP47, VAMP2-mCherry, and Stx4-EGFP.

As the SNARE complex mediates membrane fusion between cargo vesicles and the target membrane, we next sought to determine which SNAP and syntaxin proteins are involved in AMPAR exocytosis during the early phase of LTP, which is independent of protein synthesis. Previous studies reported that after chemical induction of LTP (cLTP) in hippocampal neurons with glycine, SNAP47, Stx3 and VAMP2 are required for activity-dependent AMPAR exocytosis ^10^, as determined by measuring surface levels of the AMPAR subunit GluA1 20–25 min after glycine washout—the expression phase of LTP. Additionally, SNAP25, Stx1 and VAMP2 contribute to constitutive AMPAR exocytosis ^10,28^. To determine whether SNAP47 mediates rapid exocytic trafficking of AMPAR vesicles immediately following LTP induction, we overexpressed dominant negative (DN) mutants of SNAP family members ^29,30^ in mouse hippocampal neurons in dissociated culture, chemically induced LTP and fixed cells 5 minutes after glycine washout for immunofluorescence staining of GluA1 on the cell surface. Image analysis revealed that, expression of either the SNAP29 or SNAP47 DN mutant abolished glycine-induced rapid increases in surface expression of GluA1 (Fig. 1B, C), indicating that these proteins function in activity-dependent AMPAR delivery to the PM during the early phase of LTP.

Previous studies have shown that the t-SNARE Stx3 is required for AMPAR exocytosis during LTP ^10^, whereas other studies demonstrated that at CA3-CA1 synapses, knockout of syntaxin-4 (Stx4) impairs both basal transmission and LTP ^31,32^. To determine which syntaxin functions in activity-dependent AMPAR exocytosis upon LTP induction, we depleted Stx3 or Stx4 by shRNA-mediated gene silencing (Fig. S1) in hippocampal neurons. Immunostaining and confocal microscopy revealed that depletion of Stx3, not Stx4, inhibited glycine-induced surface expression of GluA1 (Fig. 1D, E). Moreover, in dendrites of mouse hippocampal neurons, the AMPARs colocalized with VAMP2 on vesicular structures (Fig. 1F). Together, these data indicate that the SNARE proteins SNAP29, SNAP47 and Stx3 are implicated in activity-dependent AMPAR exocytic trafficking during the early phase of LTP.

Among SNAP family members, SNAP47 is not only highly expressed in the brain and distributed to both pre- and postsynaptic compartments of glutamatergic synapses ^33,34^, but is also unique in possessing an evolutionarily conserved pleckstrin homology (PH)-like domain with the potential to bind phosphoinositides. As PM PI4P is required for AMPAR exocytosis ^25^, we focused on the role of SNAP47 in LTP stimulus-induced, PI4P-dependent AMPAR PM trafficking in this study. To this end, we first determined whether SNAP47 forms complexes with VAMP2 and Stx3 by co-immunoprecipitation (coIP) assays of ectopically expressed proteins in HEK293T cells. In line with previous findings that SNARE interactions are promiscuous ^35,36^, similar to SNAP23, SNAP47 interacted with Stx3 and VAMP2, and could also complex with Stx4 (Fig. 1G–I). These data together prompted us to investigate whether and how PM PI4P regulates AMPAR exocytosis mediated by the SNAP47–Stx3–VAMP2 SNARE complex.

### SNAP47 binds phosphoinositides via the PH-like domain

As mentioned above, SNAP47 contains an evolutionarily conserved PH-like domain with the potential to bind PIPs (Fig. 2A), distinguishing it from other SNAP family members. Indeed, lipid blot overlay assays with recombinant proteins showed that SNAP47 recognized all seven PIP species, with preferential binding to PI4P, PI(3,5)P_2_ and PI(4,5)P_2_ (Fig. 2B, C). In contrast, neither Stx3, an integral membrane protein, nor endophilin A1 (EndoA1)—which binds to membranes via its membrane curvature-sensing N-BAR domain ^37^—interacted with PIPs on lipid blots (Fig. 2B,C). Moreover, deletion of the PH-like domain abrogated the PIP-binding capacity of SNAP47 (Fig. 2B, C).

**Fig. 2.**
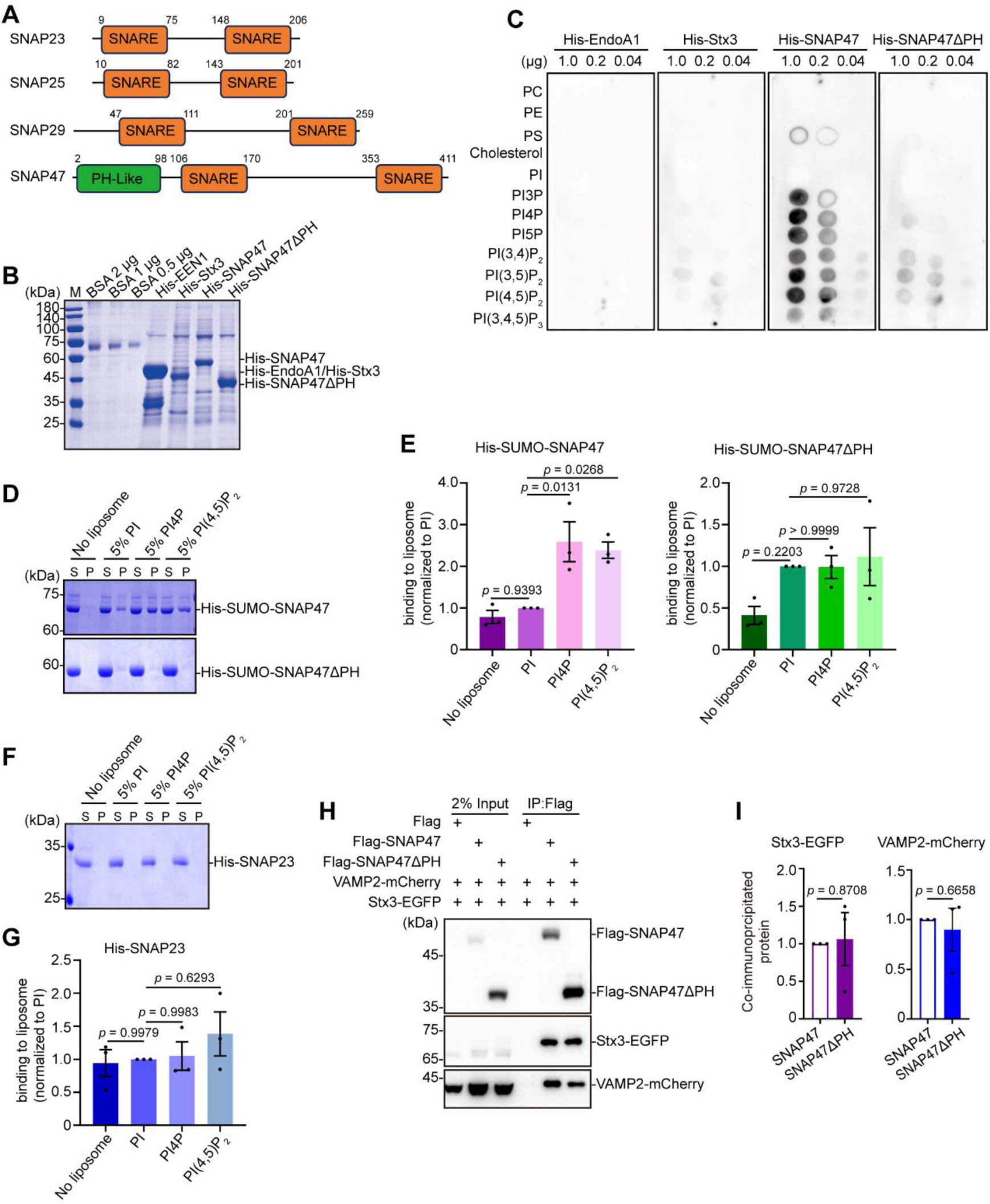
SNAP47 binds to phosphoinositides via the PH-like domain. (A) Schematic illustration of the domain structures of SNAP family proteins. (B) His-tagged SNAP47, SNAP47△PH, Stx3, and EndoA1 were expressed in *E. coli* and purified with Ni-NTA agarose. (C) Purified recombinant proteins were incubated with nitrocellulose membranes spotted with varying amounts of lipids and subsequently immunoblotted with anti-His antibodies. (D) Representative co-sedimentation assay showing the binding of His-SUMO-SNAP47 or His-SUMO-SNAP47ΔPH with liposomes. Protein-liposome mixtures were separated by centrifugation into supernatants (S) and pellets (P), and analyzed by SDS‒PAGE and Coomassie blue staining. (E) Quantification of band intensities from the co-sedimentation assay shown in (D). Data represent mean ± SEM, n = 3 independent experiments. Statistical significance was determined by one-way ANOVA analysis of variance with a Tukey post hoc test. (F) Representative co-sedimentation assay showing the binding of His-SNAP23 with liposome. (G) Quantification of band intensities from the co-sedimentation assay shown in (F). Data represent mean ± SEM, n = 3 independent experiments. Statistical significance was determined by one-way ANOVA analysis of variance with a Tukey post hoc test. (H) HEK293T cells co-transfected with constructs expressing VAMP2-mCherry, Stx3-EGFP, and Flag-SNAP47 or Flag-SNAP47ΔPH were lysed and subjected to immunoprecipitation using anti-Flag antibody-conjugated agarose beads. Input and bound fractions were analyzed by SDS‒PAGE followed by immunoblotting with antibodies against Flag, GFP, and RFP. (I) Quantification of SNAP47or SNAP47ΔPH binding to VAMP2 and Stx3 from the co-IP assays in (H). Data represent mean ± SEM, n = 3 independent experiments. Statistical significance was determined by two-tailed Student’s t-test.

Next we performed liposome sedimentation assays to corroborate the PIP-binding capacity of SNAP47. Compared with SNAP23, binding of SNAP47 to liposomes increased significantly in the presence of either PI4P or PI(4,5)P_2_, but not PI, and the protein–liposome association was severely impaired upon removal of the PH-like domain (Fig. 2D–G). Collectively, these data indicate that SNAP47 binds PI4P and PI(4,5)P_2_ via its PH-like domain. Notably, the protein–protein interactions of the SNAP47ΔPH mutant with Stx3 and VAMP2 remained intact in coIP assays (Fig. 2H, I), indicating that the PH-like domain is specifically required for the protein–lipid interaction rather than for binding to other SNARE proteins.

### PI4P promotes membrane fusion mediated by the SNAP47–Stx3–VAMP2 SNARE complex

Through electrostatic interactions, PM PI(4,5)P_2_ potentiates exocytosis by activating CAPS, recruiting the Ca^2+^ sensor Syt-1 and Munc13, and co-clustering with syntaxin-1 (Stx1) to facilitate assembly of the membrane fusion machinery at vesicle docking sites ^20,21,24,38–40^. Given that both PI(4,5)P_2_ and PI4P are anionic phospholipids that regulate effector proteins via electrostatic interactions, we reasoned that elevated PM PI4P could promote membrane fusion by recruiting SNAP47. In our previous studies, lipidomics analysis revealed that the molar ratio of both PI(4,5)P_2_ and PI4P to PI, as well as to total phospholipids, are higher in hippocampal neurons than in neuroblastoma cells, epithelial cells, and fibroblasts ^25^. To determine PM levels of these phosphoinositides, we performed lipidomics analysis of the PM isolated by surface protein biotinylation followed by streptavidin affinity purification (Fig. 3A, B). Compared with the PM of HeLa cells, the neuronal PM exhibited distinct fatty acid side-chain profiles in PI and PIPs (Fig. 3C and Table 1; Table S1). Notably, the proportion of PI4P among all PIPs detected in the neuronal PM was substantially higher (neuron: 39.52%; HeLa: 8.35%) and, remarkably, comparable to that of PM PI(4,5)P_2_ (neuron: 38.74%; HeLa: 76.13%) (Fig. 3D and Table 1; Table S1). These results suggest that PI4P plays a direct role—rather than merely serving as a metabolic precursor for PI(4,5)P_2_—in regulating SNARE-mediated exocytic fusion in neurons.

**Fig. 3.**
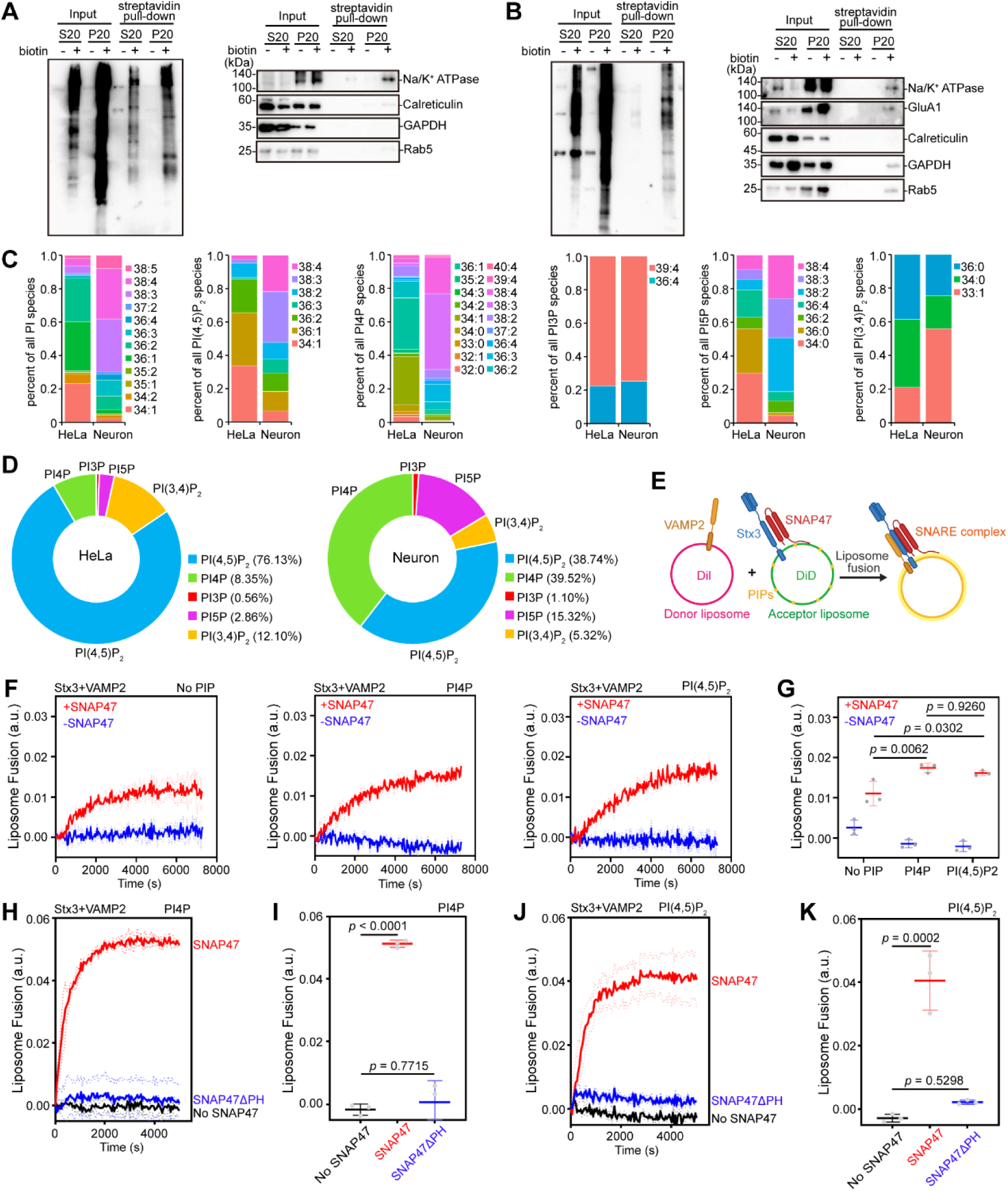
PI4P promotes membrane fusion mediated by the SNAP47–syntaxin-3–VAMP2 SNARE complex. (A) The plasma membrane of HeLa cells was isolated by surface biotinylation and streptavidin affinity purification, and subjected to lipidomics analysis. Shown are immunoblots probed with streptavidin (left) and antibodies to PM proteins (right, GAPDH served as loading control for input). (B) Same as (A) except that the PM was isolated from DIV16 rat hippocampal neurons. (C) Results from lipidomics analyses of the PMs isolated in (A) and (B). Shown are fatty acid side-chain profiles of PI and PIPs. The molar ratio of each PIP class was calculated by normalization to total PIs. n = 4 biological replicates. (D) The relative abundance of the PIP species in the PM of HeLa cells and rat hippocampal neurons. n = 4 biological replicates. (E) Schematic of the *in vitro* membrane fusion assay with reconstituted proteoliposomes. (F) *In vitro* membrane fusion measured by Förster resonance energy transfer (FRET) between donor and acceptor liposomes reconstituted without phosphoinositides, with 5% PI4P, or with 5% PI(4,5)P_2_. (G) Quantification of fusion efficiency at the end of the reactions from (F). Data represent mean ± SD, n = 3 independent experiments. Statistical significance was determined by one-way ANOVA analysis of variance with a Tukey post hoc test. (H) Membrane fusion measured by FRET between donor liposomes and acceptor liposomes reconstituted with 5% PI4P, using wild-type SNAP47 or its mutant SNAP47ΔPH. (I) Quantification of fusion efficiency at the end of the reaction from (H). Data represent mean ± SD, n = 3 independent experiments. Statistical significance was determined by one-way ANOVA analysis of variance with a Tukey post hoc test. (J) Membrane fusion measured by FRET between donor liposomes and acceptor liposomes reconstituted with 5% PI(4,5)P_2_, using wild-type SNAP47 or its mutant SNAP47ΔPH. (K) Quantification of fusion efficiency at the end of the reaction from (J). Data represent mean ± SD, n = 3 independent experiments. Statistical significance was determined by one-way ANOVA analysis of variance with a Tukey post hoc test.

**Table 1.** Comparison of plasma membrane PIP profiles between HeLa cells and rat hippocampal neurons.

| **Composition** | **HeLa**<br>**N = 4<sup>1</sup>** | **Neuron**<br>**N = 4<sup>1</sup>** | **P-value** |
| --- | --- | --- | --- |
| PI(3)P | 0.52 (0.47, 0.65) | 1.13 (0.95, 1.25) | 0.029<sup>2</sup> |
| PI(3)P-36:4 | 20.05 (17.26, 27.29) | 22.92 (20.86, 29.82) | 0.343<sup>2</sup> |
| PI(3)P-39:4 | 79.95 (72.71, 82.74) | 77.08 (70.18, 79.14) | 0.343<sup>2</sup> |
| PI(3,4)P<sub>2</sub> | 11.45 (10.80, 13.40) | 4.96 (4.42, 6.23) | 0.029<sup>2</sup> |
| PI(3,4)P<sub>2</sub>-33:1 | 21.25 (15.59, 26.87) | 55.18 (48.34, 63.44) | 0.029<sup>2</sup> |
| PI(3,4)P<sub>2</sub>-34:0 | 40.86 (36.85, 43.61) | 21.03 (15.21, 24.22) | 0.029<sup>2</sup> |
| PI(3,4)P<sub>2</sub>-36:0 | 37.77 (33.52, 43.56) | 26.41 (18.55, 30.24) | 0.029<sup>2</sup> |
| PI(4)P | 8.35 (7.72, 8.98) | 40.32 (35.82, 43.21) | 0.029<sup>2</sup> |
| PI(4)P-32:0 | 3.51 (3.07, 3.91) | 0.14 (0.12, 0.22) | 0.029<sup>2</sup> |
| PI(4)P-32:1 | 1.43 (1.24, 1.64) | 0.37 (0.33, 0.38) | 0.029<sup>2</sup> |
| PI(4)P-33:0 | 1.56 (1.25, 1.88) | 0.37 (0.27, 0.43) | 0.029<sup>2</sup> |
| PI(4)P-34:0 | 3.83 (3.67, 4.31) | 0.26 (0.21, 0.28) | 0.029<sup>2</sup> |
| PI(4)P-34:1 | 28.96 (28.44, 29.34) | 2.47 (2.42, 2.65) | 0.029<sup>2</sup> |
| PI(4)P-34:2 | 1.67 (1.52, 2.03) | 0.93 (0.84, 0.99) | 0.029<sup>2</sup> |
| PI(4)P-34:3 | 0.44 (0.38, 0.48) | 0.41 (0.36, 0.44) | 0.486<sup>2</sup> |
| PI(4)P-35:2 | 1.76 (1.58, 2.22) | 0.91 (0.87, 1.05) | 0.029<sup>2</sup> |
| PI(4)P-36:1 | 30.77 (30.09, 31.50) | 1.75 (1.50, 2.06) | 0.029<sup>2</sup> |
| PI(4)P-36:2 | 9.69 (9.28, 10.42) | 4.64 (4.39, 4.83) | 0.029<sup>2</sup> |
| PI(4)P-36:3 | 0.86 (0.77, 1.02) | 10.85 (9.99, 11.20) | 0.029<sup>2</sup> |
| PI(4)P-36:4 | 1.41 (1.20, 1.57) | 1.93 (1.86, 2.12) | 0.029<sup>2</sup> |
| PI(4)P-37:2 | 0.73 (0.50, 0.96) | 1.39 (1.29, 1.44) | 0.029<sup>2</sup> |
| PI(4)P-38:2 | 6.03 (5.61, 6.91) | 5.35 (5.02, 5.46) | 0.057<sup>2</sup> |

| **Composition** | **HeLa**<br>N = 4<sup>1</sup> | **Neuron**<br>N = 4<sup>1</sup> | **P-value** |
| --- | --- | --- | --- |
| PI(4)P-38:3 | 1.79 (1.70, 1.97) | 44.21 (43.81, 46.58) | 0.029<sup>2</sup> |
| PI(4)P-38:4 | 2.77 (2.44, 3.43) | 21.91 (20.80, 22.88) | 0.029<sup>2</sup> |
| PI(4)P-39:4 | 1.47 (1.40, 1.56) | 0.61 (0.53, 0.90) | 0.029<sup>2</sup> |
| PI(4)P-40:4 | 0.35 (0.16, 0.55) | 0.73 (0.60, 0.87) | 0.057<sup>2</sup> |
| PI(4,5)P<sub>2</sub> | 75.60 (75.33, 76.92) | 38.82 (34.35, 43.12) | 0.029<sup>2</sup> |
| PI(4,5)P<sub>2</sub>-34:1 | 33.54 (32.88, 34.84) | 7.54 (5.07, 8.56) | 0.029<sup>2</sup> |
| PI(4,5)P<sub>2</sub>-36:1 | 32.05 (30.86, 32.61) | 10.71 (5.30, 17.94) | 0.029<sup>2</sup> |
| PI(4,5)P<sub>2</sub>-36:2 | 20.14 (19.12, 20.95) | 9.90 (9.37, 12.76) | 0.029<sup>2</sup> |
| PI(4,5)P<sub>2</sub>-36:3 | 1.00 (0.93, 1.13) | 8.27 (7.30, 9.39) | 0.029<sup>2</sup> |
| PI(4,5)P<sub>2</sub>-38:2 | 8.91 (8.00, 9.53) | 9.84 (9.22, 10.56) | 0.343<sup>2</sup> |
| PI(4,5)P<sub>2</sub>-38:3 | 2.10 (1.85, 2.30) | 31.20 (23.98, 36.87) | 0.029<sup>2</sup> |
| PI(4,5)P<sub>2</sub>-38:4 | 2.57 (2.29, 2.72) | 21.75 (19.40, 24.27) | 0.029<sup>2</sup> |
| PI(5)P | 2.95 (2.42, 3.31) | 15.36 (14.03, 16.61) | 0.029<sup>2</sup> |
| PI(5)P-34:0 | 29.08 (25.16, 34.22) | 4.82 (3.93, 5.22) | 0.029<sup>2</sup> |
| PI(5)P-36:0 | 26.54 (23.39, 29.80) | 1.63 (1.34, 2.03) | 0.029<sup>2</sup> |
| PI(5)P-36:2 | 7.10 (5.63, 7.85) | 6.71 (5.73, 8.29) | 0.886<sup>2</sup> |
| PI(5)P-36:4 | 17.22 (14.57, 17.58) | 5.74 (4.75, 6.30) | 0.029<sup>2</sup> |
| PI(5)P-38:2 | 6.95 (3.84, 8.77) | 32.28 (29.58, 34.16) | 0.029<sup>2</sup> |
| PI(5)P-38:3 | 5.78 (4.94, 6.84) | 22.88 (22.56, 24.71) | 0.029<sup>2</sup> |
| PI(5)P-38:4 | 7.30 (6.10, 11.32) | 24.43 (23.68, 27.72) | 0.029<sup>2</sup> |

<sup>1</sup>Median (Q1, Q3)
<sup>2</sup>Wilcoxon rank sum exact test

*In vitro* assembly of the SNARE complex catalyzes fusion of proteoliposomes. As SNAP47 forms complexes with both Stx3 and VAMP2 and interacts with PI4P, we next investigated the function of PI4P in membrane fusion using *in vitro* assays reconstituted with liposomes and recombinant SNARE proteins (Fig. 3E) ^41,42^. Indeed, similar to PI(4,5)P_2_, PI4P promoted lipid mixing between donor proteoliposomes containing VAMP2 and acceptor proteoliposomes containing Stx3 in the presence of SNAP47 (Fig. 3F, G), indicating that PI4P can facilitate membrane fusion mediated by the SNAP47–Stx3–VAMP2 SNARE complex. Moreover, for both PI4P and PI(4,5)P_2_, the fusion-promoting effect was abolished when wild-type SNAP47 was substituted with the ΔPH mutant (Fig. 3H–K). Together, these data demonstrate that PI4P is as efficient as PI(4,5)P_2_ in promoting membrane fusion mediated by SNARE complexes containing the PIP-binding SNAP47.

### PM PI4P promotes the assembly of the SNARE complex

Having established that PI4P promotes the SNAP47–Stx3–VAMP2 SNARE complex-mediated membrane fusion *in vitro*, we next asked whether PI4P enhances SNARE complex assembly in cells. To address this, we manipulated PM PI4P levels in HEK293T cells co-expressing the SNARE proteins and then performed coIP assays. Overexpression of PI4KIIIα, which increases PM PI4P ^25^, enhanced the SNAP47–VAMP2 interaction, but did not affect the SNAP47–Stx3 or VAMP2–Stx3 interactions (Fig. 4A–F). To corroborate that PI4P strengthens the SNAP47–VAMP2 interaction, we conducted *in vitro* protein-binding assays by incubating bacterially expressed recombinant VAMP2 immobilized on glutathione agarose beads with His-tagged SNAP47 and Stx3. Compared with PI (as negative control), addition of PI4P but not PI(4,5)P_2_ enhanced SNAP47 binding to VAMP2 (Fig. 4G, H). Moreover, deletion of SNAP47’s PH-like domain weakened its interaction with VAMP2 in the presence of PI4P (Fig. 4I, J), suggesting that PI4P-enhanced SNAP47–VAMP2 interaction requires the PH-like domain. Notably, neither PI4P nor PI(4,5)P_2_ affected VAMP2 binding to Stx3, regardless of whether SNAP47 was wild-type or deleted of the PH-like domain (Fig. 4G–J). Further, in coIP assays, treating cells co-expressing the SNAREs with GSK-A1, an inhibitor of PI4KIIIα that depletes PM PI4P but not PI(4,5)P_2_ ^25^, weakened the interaction of VAMP2 with SNAP47 but not SNAP23 (Fig. 4K, L). Notably, GSK-A1 treatment did not affect SNAP47–Stx3 or SNAP23–Stx3 interactions (Fig. 4K, L). Collectively, these data indicate that PI4P specifically enhances the SNAP47–VAMP2 interaction in a PH-like domain-dependent manner, which may facilitate the assembly of the SNAP47–Stx3–VAMP2 SNARE complex.

**Fig. 4.**
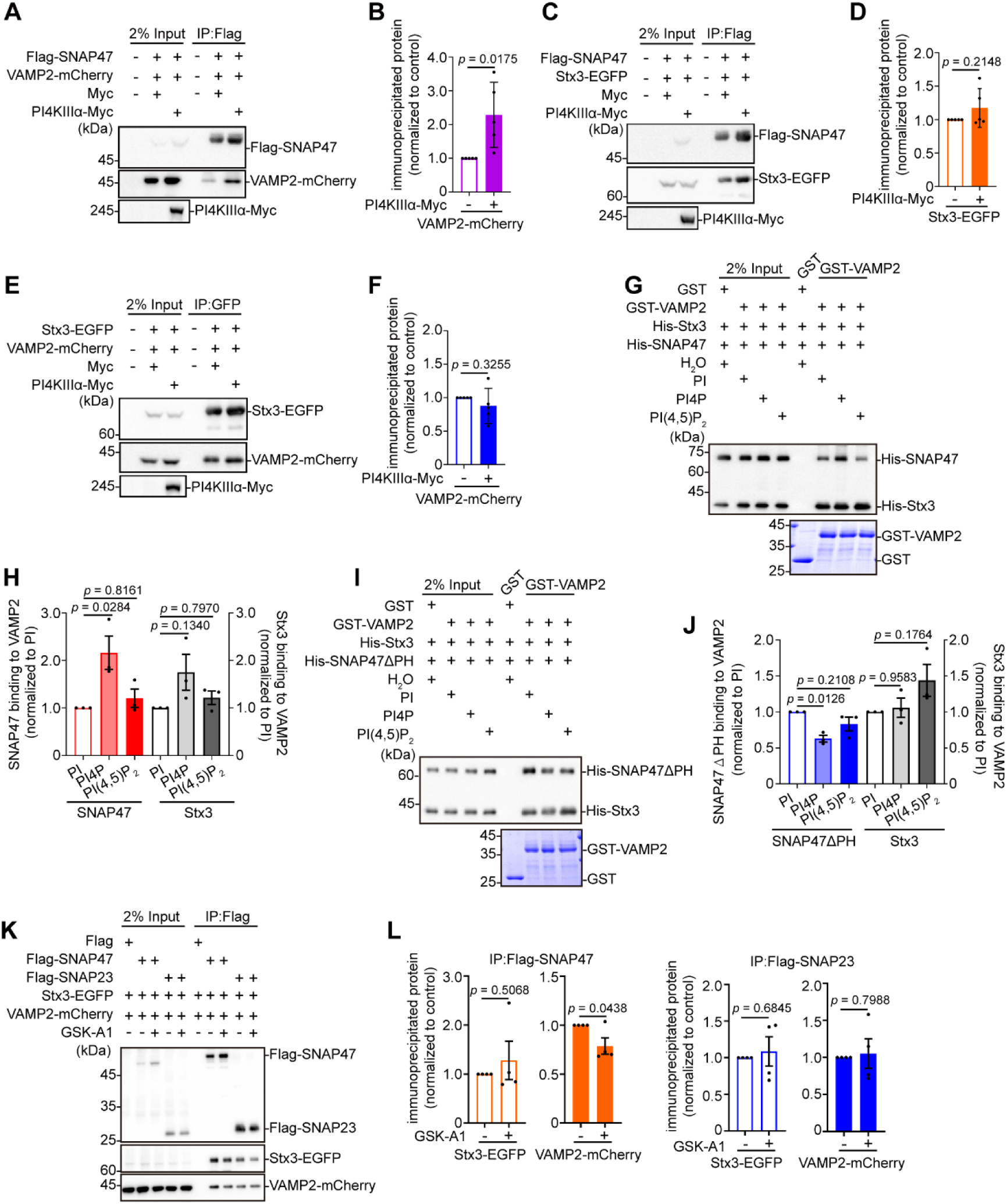
PI4P promotes the assembly of the SNAP47–syntaxin-3–VAMP2 SNARE complex. (A) HEK293T cells co-transfected with constructs expressing Flag-SNAP47, VAMP2-mCherry, and PI4KIIIα-Myc were lysed and subjected to IP using anti-Flag-conjugated agarose beads. Input and bound fractions were analyzed by SDS‒PAGE followed by immunoblotting with antibodies against Flag, RFP, and Myc. (B) Quantification of SNAP47‒VAMP2 interaction in (A). Data represent mean ± SEM, n = 5 independent experiments. Statistical significance was determined by two-tailed Student’s t-test. (C) HEK293T cells co-transfected with constructs expressing Flag-SNAP47, Stx3-EGFP, and PI4KIIIα-Myc were lysed and subjected to IP using anti-Flag-conjugated agarose beads. (D) Quantification of SNAP47‒Stx3 interaction in (C). Data represent mean ± SEM, n = 5 independent experiments. Statistical significance was determined by two-tailed Student’s t-test. (E) HEK293T cells co-transfected with constructs expressing Stx3-EGFP, VAMP2-mCherry, and PI4KⅢα-Myc were lysed and subjected to IP using anti-GFP-conjugated agarose beads. (F) Quantification of VAMP2‒Stx3 interaction in (E). Data represent mean ± SEM, n = 5 independent experiments. Statistical significance was determined by two-tailed Student’s t-test. (G) GST-tagged VAMP2 immobilized to glutathione-Sepharose was incubated with His-SNAP47 and His-Stx3 in the presence of PI, PI4P, or PI(4,5)P_2_. (H) Quantification of VAMP2‒SNAP47 and VAMP2‒Stx3 interactions in (G). Data represent mean ± SEM, n = 3 independent experiments. Statistical significance was determined by one-way ANOVA analysis of variance with a Tukey post hoc test. (I) GST-tagged VAMP2 immobilized to glutathione-Sepharose was incubated with His-SNAP47ΔPH and His-Stx3 in the presence of PI, PI4P, or PI(4,5)P_2_. (J) Quantification of VAMP2‒SNAP47ΔPH and VAMP2‒Stx3 interactions in (I). Data represent mean ± SEM, n = 3 independent experiments. Statistical significance was determined by one-way ANOVA analysis of variance with a Tukey post hoc test. (K) HEK293T cells co-transfected with constructs expressing VAMP2-mCherry, Stx3-EGFP, and either Flag-SNAP47 or Flag-SNAP23 were treated with the PI4KIIIα inhibitor GSK-A1, lysed and subjected to immunoprecipitation with anti-Flag-conjugated agarose beads. (L) Quantification of SNARE complex assembly in (K). Data represent mean ± SEM, n = 4 independent experiments. Statistical significance was determined by two-tailed Student’s t-test.

### SNAP47 is recruited to PI4P-enriched PM microdomains to mediate AMPAR exocytic fusion during LTP

Having demonstrated that the SNAP47–PI4P interaction promotes SNARE-mediated membrane fusion, we next investigated whether this protein–lipid interaction contributes to activity-dependent AMPAR exocytosis in neurons. SNARE proteins have been reported to concentrate in PM microdomains that may serve as signposts for fusion sites for exocytic vesicles ^43,44^. Notably, unlike SNAP25, which associates with the PM through palmitoylation ^45^, SNAP47 lacks identifiable membrane anchor domains ^33^. To determine whether SNAP47 is recruited to the PM via its interaction with PI4P, we examined changes in SNAP47 signals at the PM during LTP induction using TIRF live imaging. Glycine application induced a rapid increase in wild-type, but not ΔPH mutant SNAP47 signals at the PM (Fig. 5A and Video 2), and this recruitment was inhibited by the PI4KIIIα inhibitor GSK-A1, suggesting that SNAP47 localizes to the PM through PI4P binding (Fig. 5B and Video 3). We further observed SNAP47 recruitment to dendritic PM regions with elevated PI4P levels as detected by EGFP-PH^OSBP^ (Fig. 5C and Video 4). In contrast, SNAP47 recruitment sites did not overlap with PM ‒signals of PH^PLCδ1^-EGFP, the PI(4,5)P_2_ probe (Fig. 5D and Video 4). Collectively, these data indicate that SNAP47 is recruited to PI4P-enriched microdomains in the dendritic PM of potentiated neurons. Next, to determine whether Stx3, the transmembrane t-SNARE, is involved in activity-induced PM recruitment of SNAP47, we depleted Stx3 by shRNA-mediated knockdown (KD) (Fig. S2A). Silencing Stx3 had no effect on glycine-induced SNAP47 accumulation at the dendritic PM (Fig. 5E and Video 5), indicating that activity-triggered SNAP47 PM localization is PI4P-dependent and Stx3-independent.

**Fig. 5.**
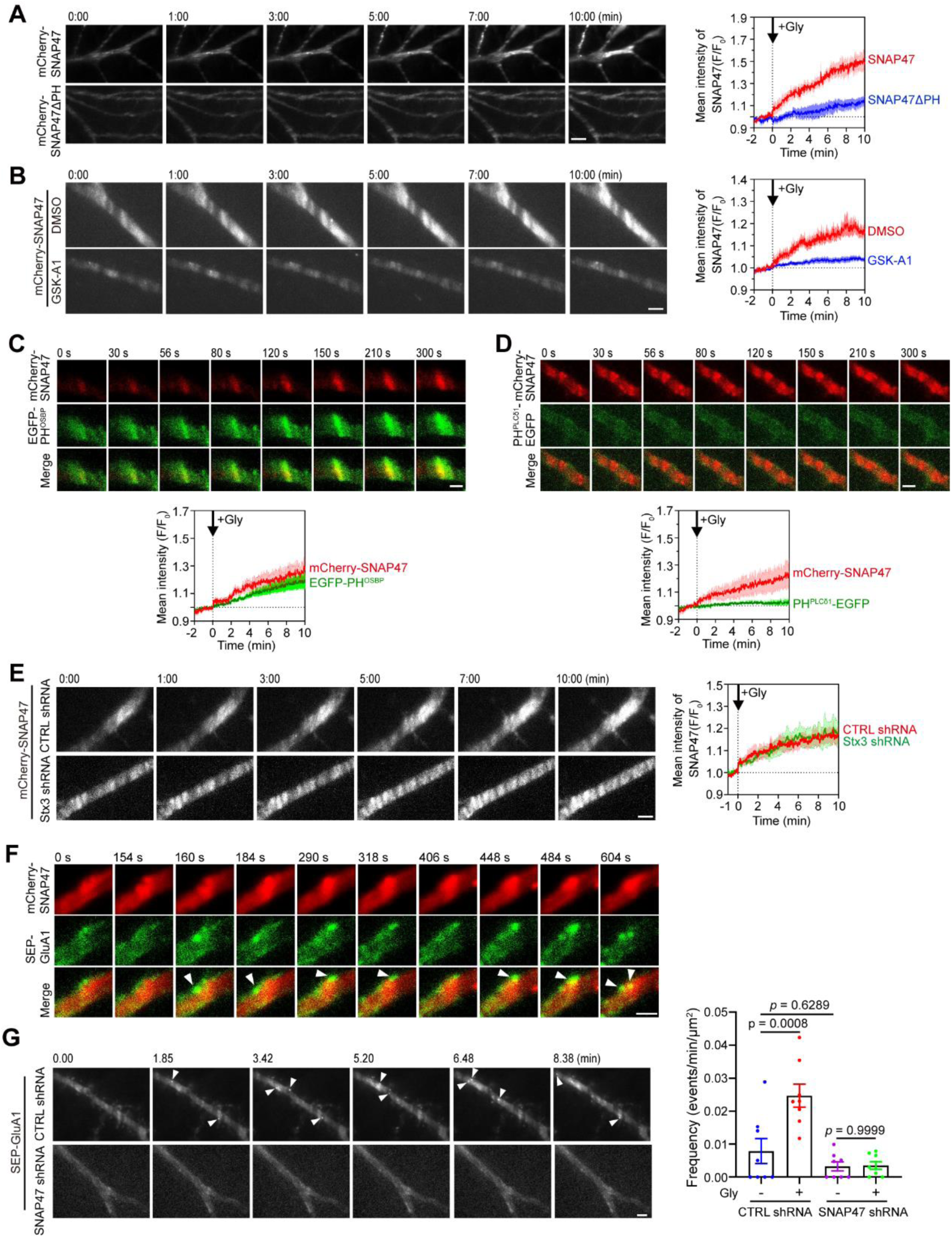
Plasma membrane PI4P recruits SNAP47 and promotes exocytosis of AMPAR vesicles. (A) Cultured rat hippocampal neurons were transfected on DIV11 with constructs expressing either mCherry-SNAP47 or mCherry-SNAP47ΔPH and imaged live by TIRF microscopy on DIV16-18 before and after glycine application. Shown are representative images (left) and normalized kinetics of fluorescence intensity changes (right). Data represent mean ± SEM (mCherry-SNAP47, n = 5 neurons; mCherry-SNAP47ΔPH, n = 6 neurons). Scale bar, 3 μm. (B) Cultured rat hippocampal neurons expressing mCherry-SNAP47 were imaged live by TIRF microscopy before and after glycine application. Neurons were treated with DMSO (vehicle control) or the PI4KIIIα inhibitor GSK-A1 during cLTP induction. Data represent mean ± SEM (DMSO, n = 6 neurons; GSK-A1, n = 6 neurons). Scale bar, 3 μm. (C) Cultured rat hippocampal neurons co-expressing mCherry-SNAP47 and EGFP-PH^OSBP^ were imaged live by TIRF microscopy before and after glycine application. Data represent mean ± SEM, n = 4 neurons. Scale bar, 2 μm. (D) Same as (C) except that neurons were co-transfected with constructs expressing mCherry-SNAP47 and PH^PLCδ1^-EGFP. Data present mean ± SEM, n = 4 neurons. Scale bar, 2 μm. (E) Cultured rat hippocampal neurons were co-transfected with constructs expressing mCherry-SNAP47 and non-targeting control shRNA (CTRL shRNA) or Stx3-targeting shRNA (Stx3 shRNA), and imaged live by TIRF microscopy before and after glycine application. Data represent mean ± SEM (CTRL shRNA, n = 6 neurons; Stx3 shRNA, n = 5 neurons). Scale bar, 3 μm. (F) Cultured rat hippocampal neurons co-expressing mCherry-SNAP47 and SEP-GluA1 were imaged live by TIRF microscopy before and after glycine application. Shown are representative images of glycine-stimulated neurons. Arrowheads indicate SEP-GluA1 exocytosis at the plasma membrane with enhanced mCherry-SNAP47 signals. Scale bar, 2 μm. (G) Cultured rat hippocampal neurons were co-transfected with constructs expressing SEP-GluA1 and non-targeting control shRNA (CTRL shRNA) or SNAP47-targeting shRNA (SNAP47 shRNA), and imaged live by TIRF microscopy before and after glycine application. Shown are representative images (left) and quantification of AMPAR exocytic events (right). Arrowheads indicate SEP-GluA1 exocytosis. Data represent mean ± SEM. Scale bar, 3 μm.

As SNAP47 is required for activity-induced AMPAR surface expression, we next examined the spatiotemporal relationship between AMPAR exocytosis and SNAP47 PM recruitment. TIRF imaging revealed that, following glycine application, SEP-GluA1 was repeatedly exocytosed at SNAP47-enriched dendritic PM regions (Fig. 5F and Video 6), suggesting that SNAP47-clustered PM microdomains serve as hotspots for membrane fusion. Finally, we determined whether activity-dependent AMPAR vesicle exocytosis requires SNAP47. Indeed, silencing SNAP47 expression (Fig. S2B) inhibited glycine-induced AMPAR exocytosis in hippocampal neurons (Fig. 5G and Video 7). All together, these data demonstrate that locally enriched PI4P facilitates SNARE-mediated fusion between AMPAR transport vesicles and the PM.

### The SNAP47–PI4P interaction is required for activity-dependent synaptic expression of AMPARs, LTP, and long-term memory

Having established that SNAP47 is required for activity-dependent AMPAR exocytosis at PI4P-enriched PM microdomains, we next sought to verify the mechanistic role of the SNAP47–PI4P interaction in synaptic delivery of AMPARs and LTP. First, we performed SNAP47 gene silencing and rescue (Fig. S2C, D) experiments in cultured hippocampal neurons. Consistent with the inhibitory effect of the dominant negative mutant, postsynaptic knockdown of SNAP47 completely abolished activity-dependent surface insertion and synaptic expression of AMPARs following LTP induction (Fig. 6A–D). Moreover, wild-type SNAP47, but not the ΔPH mutant, rescued the AMPAR trafficking phenotypes in KD neurons (Fig. 6A–D).

**Fig. 6.**
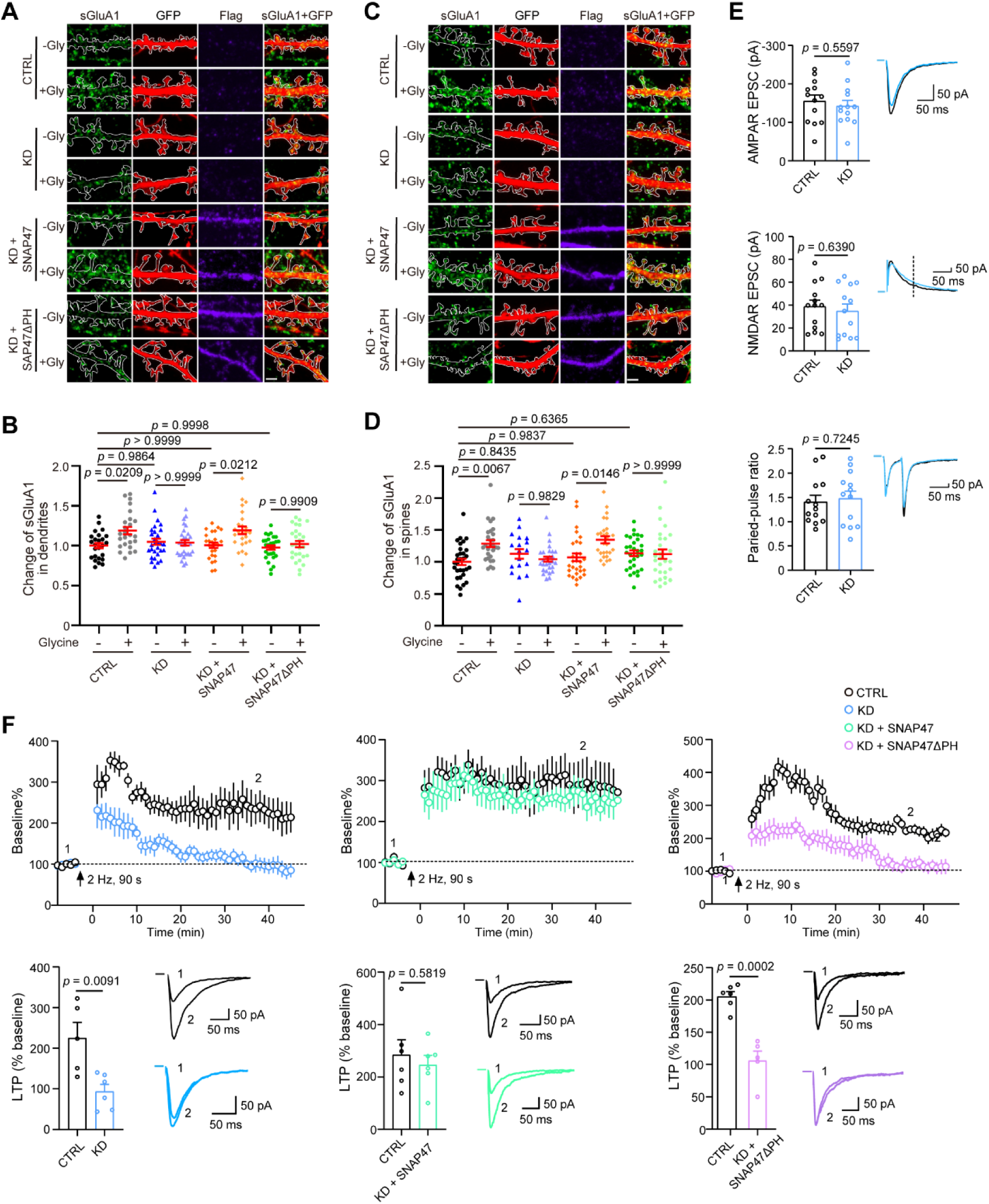
The SNAP47‒PI4P protein‒lipid interaction is required for activity-dependent AMPAR exocytosis, LTP expression and long-term memory. (A) Cultured mouse hippocampal neurons were co-transfected on DIV11 with the Flag vector and construct co-expressing EGFP and non-targeting control shRNA (CTRL) or SNAP47-targeting shRNA (KD), or constructs for SNAP47 shRNA and Flag-tagged SNAP47 or SNAP47ΔPH. On DIV16, cLTP was induced and neurons were fixed 5 min following glycine washout and immunostained for GFP, Flag, and surface-expressed GluA1. Shown are representative confocal fluorescence images. Scale bar, 2 μm. (B) Quantification of surface GluA1 in dendrites in (A). Data represent mean ± SEM, n = 25-29 neurons from two independent experiments, 1-3 dendrites/neuron. Statistical significance was determined by one-way ANOVA analysis of variance with a Tukey post hoc test. (C) Same as (A) except that cells were fixed 30 min following glycine washout. Shown are representative confocal fluorescence images. (D) Quantification of surface GluA1 in dendritic spines in (C). Data represent mean ± SEM, n = 18-31 neurons from two independent experiments, 1-3 dendrites/neurons. Statistical significance was determined by one-way ANOVA analysis of variance with a Tukey post hoc test. Scale bar, 2 μm. (E) AAVs expressing SNAP47 shRNA were stereotaxically delivered into the hippocampal CA1 region of mouse brain at postnatal day 0 (P0). Acute hippocampal slices were subsequently prepared between P14 and P21 for electrophysiological characterization of CA1 pyramidal neurons. Bar graphs summarize the amplitudes of AMPAR-mediated excitatory postsynaptic currents (AMPAR-EPSCs, top), NMDAR-mediated excitatory postsynaptic currents (NMDAR-EPSCs, center), and the paired-pulse ratio in control (CTRL, uninfected) and AAV-infected (KD) CA1 pyramidal neurons (bottom). Data are presented as mean ± SEM: AMPAR-EPSCs (CTRL, −155.19 ± 15.63 pA; SNAP47 KD, - 142.50 ± 14.71 pA; n = 13 neurons from 3 mice), NMDAR-EPSCs (CTRL, 38.79 ± 5.58 pA; SNAP47 KD, 34.87 ± 6.07 pA; n = 13 neurons from 3 mice), and paired-pulse ratio (CTRL, 1.407 ± 0.134; SNAP47 KD, 1.479 ± 0.152; n = 13 neurons from 3 mice). Statistical comparisons were performed using two-tailed Student’s t-tests. Representative traces from control (CTRL, black) and AAV-infected (KD, blue) neurons are shown in the inset. Scale bars, 50 pA and 50 ms. (F) Rescue of long-term potentiation (LTP) deficits in SNAP47 knockdown (KD) neurons with wild-type SNAP47, but not the SNAP47ΔPH mutant. AAVs expressing SNAP47 shRNA alone or in combination with AAVs expressing either mCherry-SNAP47 or mCherry-SNAP47ΔPH were stereotaxically delivered into the hippocampal CA1 region of mouse brain at P0. LTP analysis was performed on acute hippocampal slices prepared between P14 and P21. Time-course plots depict normalized AMPAR-EPSC amplitudes (mean ± SEM) for non-infected control neurons (CTRL, black) and AAV-infected neurons (KD, blue; KD + SNAP47 WT, green; KD + SNAP47ΔPH, purple), with values normalized to the baseline AMPAR-EPSC amplitude prior to LTP induction (arrow). Bar graphs below summarize the percentage of baseline responses measured at 35 – 45 min. Statistical comparisons include: CTRL vs KD (CTRL, n = 5; KD, n = 6; 5 mice), CTRL vs KD + SNAP47 WT (CTRL, n = 6; KD + SNAP47 WT, n = 6; 6 mice), and CTRL vs KD + SNAP47ΔPH (CTRL, n = 6; KD + SNAP47ΔPH, n = 5; 6 mice). All statistical analyses were conducted using two-tailed Student’s t-tests. Representative AMPAR-EPSC traces recorded before (1) and after (2) LTP induction from control and AAV-infected neurons are shown beneath the graphs. Scale bars, 50 pA and 50 ms.

Next, we tested the functional significance of the SNAP47–PI4P interaction in LTP by electrophysiological analysis. We introduced adeno-associated virus (AAV) encoding SNAP47-targeting shRNA into the hippocampal CA1 region of newborn mice on postnatal day 0 (P0), and performed whole-cell patch-clamp recordings of evoked excitatory postsynaptic currents (eEPSCs) in CA1 pyramidal neurons from acute brain slices on postnatal days 14–21. Silencing SNAP47 did not affect AMPAR- or NMDAR-mediated synaptic transmission, nor the paired-pulse ratio of AMPAR eEPSCs (Figure 6E). However, suppression of SNAP47 expression significantly impaired LTP in CA1 neurons following Schaffer collateral stimulation, which was restored by co-expression of RNAi-resistant wild-type SNAP47 but not the ΔPH mutant (Fig. 6F and S3A).

Finally, we determined the physiological relevance of the SNAP47–PI4P interaction by examining the effects of SNAP47 gene silencing on hippocampus-dependent learning and memory. We injected shRNA-expressing AAV into the hippocampal CA1 region of adult mice (Fig. S3B). Suppressing SNAP47 expression did not alter locomotor activity (open field and rotarod assays), anxiety levels (elevated plus maze), working memory (Y maze), or—consistent with hippocampal CA1 deletion of Stx3 ^46^—memory in contextual fear conditioning (Fig. 7A–E). However, mice injected with SNAP47 shRNA-expressing AAV exhibited deficits in longterm memory in the Morris water maze test, which were rescued by wild-type SNAP47 but not the ΔPH mutant (Fig. 7F). Collectively, these data indicate that the direct interaction between SNAP47 and PM PI4P plays a critical role in activity-induced AMPAR synaptic expression, LTP, and long-term memory.

**Fig. 7.**
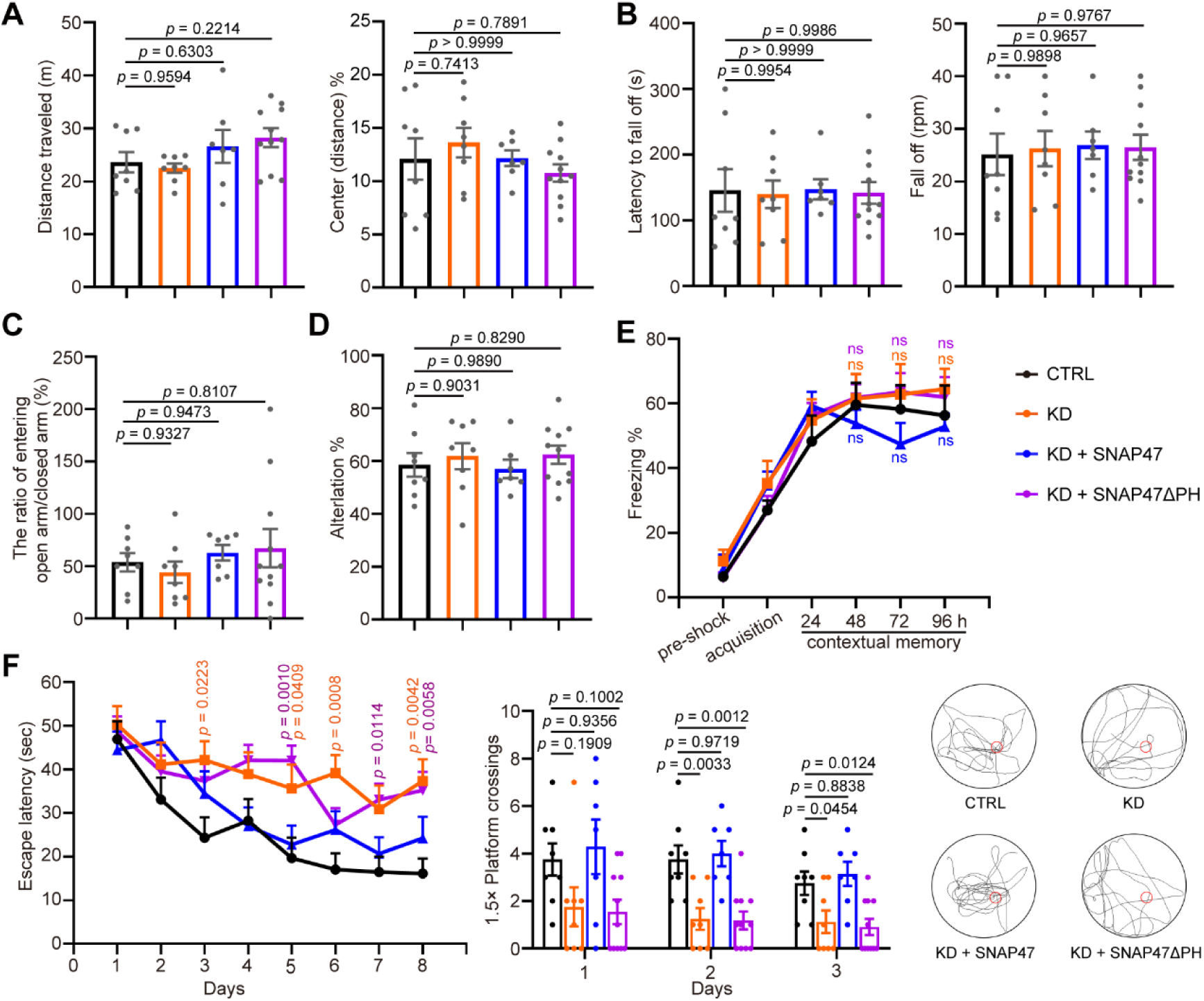
The SNAP47–PI4P protein–lipid interaction is required for long-term memory. (A–F) Eight-week-old male mice were stereotaxically injected with AAVs (CTRL shRNA + mCherry vector (CTRL), SNAP47 shRNA + mCherry vector (KD), SNAP47 shRNA + mCherry-SNAP47 (KD + SNAP47), or SNAP47 shRNA + mCherry-SNAP47ΔPH (KD + SNAP47ΔPH)). Two weeks after virus injection, mice were subjected to behavior tests: the open field (A), rotarod (B), elevated plus maze (C), Y maze (D), contextual fear conditioning (E), and the Morris water maze (F). Shown in (F) are the escape latency before escaping to the platform in the invisible platform training, number of crossings within the 1.5× platform area at the probe test, and the swim traces in the probe test. Red circles indicate positions of the submerged/invisible platform. All data represent means ± SEM for each group (CTRL, n = 8; KD, n = 8; KD + SNAP47, n = 7; KD + SNAP47ΔPH, n = 11 from two independent experiments). ns: not significant. Statistical significance was determined by one-way ANOVA analysis of variance with a Dunnett post hoc test.

## Discussion

The heterogeneous subcellular distribution of phosphoinositides and their transient changes in local concentration within organelle membranes and the PM regulate membrane trafficking and dynamics ^11^. In dendrites of potentiated hippocampal neurons, the Ca^2+^-sensing membrane tether protein E-Syt1 recruits PI4KIIIα to ER‒PM contact sites, increasing PM PI4P levels ^25,26^.

In the current study, we found that AMPAR transport vesicles undergo exocytosis at PI4P-enriched PM microdomains, and that PI4P facilitates SNARE-mediated membrane fusion by recruiting SNAP47 to the PM. These findings reveal a mechanistic role for PI4P in neuronal activity-dependent AMPAR exocytosis.

It is well established that in neuroendocrine cells, PM PI(4,5)P_2_ functions in the docking, priming and fusion of dense core granules during Ca^2+^-regulated exocytosis ^47,48^. Notably, recent studies revealed that during depolarization-induced exocytosis in neuroendocrine chromaffin cells, Ca^2+^-activated synaptojanin rapidly converts PI(4,5)P_2_ to PI4P, and elevated PI4P drives dynamin-mediated fission of endocytic vesicles from the PM ^49^. In central nervous system neurons, PI(4,5)P_2_ plays essential roles in synaptic vesicle trafficking at presynaptic terminals ^16,17,50^. Upon activity-induced exocytosis of synaptic vesicles at presynaptic sites, the Ca^2+^ sensor Syt-1 recruits PIPKIγ to catalyze local synthesis of PI(4,5)P_2_, promoting endocytic retrieval of synaptic vesicle membranes ^16^. However, postsynaptically, acute depletion of PM PI(4,5)P_2_ has no impact on LTP stimulus-induced AMPAR surface expression or LTP expression ^25,51^, suggesting a PI(4,5)P_2_-independent mechanism for exocytic trafficking of AMPAR transport vesicles. While *in vitro* assays indicate that PI(4,5)P_2_ regulates membrane fusion mediated by the canonical SNAP25–Stx1–VAMP2 SNARE complex ^52^, our findings reveal that besides its role as the immediate precursor of PI(4,5)P_2_, PI4P functions directly in SNARE-mediated exocytic fusion by recruiting SNAP47 to the dendritic PM and facilitating SNARE-mediated membrane fusion of AMPAR transport vesicles (model, Fig. S4). Together, these findings demonstrate the essential role of neuronal activity-dependent synthesis and signaling of phosphoinositides in synaptic transmission and plasticity.

Previous studies found that, the t-SNARE proteins, Stx1 and SNAP25, or Stx4 and SNAP23, form cholesterol-dependent clusters in the PM, increasing local SNARE concentrations for more efficient fusion ^43,53^. Stx1 undergoes homo-oligomerization to form clusters in the PM, a process regulated by its SNARE domain and the membrane lipids cholesterol and PI(4,5)P_2_ ^54,55^. It was reported that Stx1A clustering is mediated by electrostatic protein–lipid interactions between its C-terminal polybasic juxtamembrane helix and strongly anionic PI(4,5)P_2_, and that its sequestration in PI(4,5)P_2_ microdomains facilitates assembly of the SNARE membrane fusion machinery to mediate exocytosis ^23,38^. However, TIRF imaging of live insulin-secreting cells revealed that PI(4,5)P_2_ is evenly distributed in the PM, and its acute depletion inhibits exocytosis but not secretory granule docking or Stx1A clustering ^22^. Moreover, most recent studies using light-inducible clustering of Stx1A suggest that syntaxin clusters form through liquid–liquid phase separation of the SNARE domain, which serve as a reservoir from which Munc18 captures syntaxin monomers to form a syntaxin–Munc18 complex for efficient fusion ^56^. Notably, PI(4,5)P_2_ promotes vesicle priming mediated by Munc13 and CAPS ^21^, and also serves as an electrostatic catalyst for membrane fusion by lowering the hydration energy barrier ^57^. Given that PI(4,5)P_2_ is enriched in the PM, it is conceivable that through its negative charges and protein–lipid interactions, PI(4,5)P_2_ plays facilitatory roles at multiple steps of exocytosis. How PI4P, which is also negatively charged and more concentrated in the PM than in other organelles except the Golgi compartment, contributes to exocytosis in various cell types, has not been thoroughly explored.

Remarkably, lipidomics analysis, immunofluorescence staining, and high-resolution microscopic imaging with fluorescent lipid probes revealed that hippocampal neurons have a larger pool of PI4P in the PM compared with other cell types, and that the percentage of cellular PI4P relative to total phospholipids (5.82%) is even higher than that of PI(4,5)P_2_ (2.17%) ^25,58^. In the current study, lipidomics analysis verified that the relative abundance of PI4P is similar to that of PI(4,5)P_2_ in the neuronal PM (39.52% vs 38.74%). In potentiated neurons, PM PI4P levels are further increased upon LTP induction by enhanced synthesis catalyzed by PI4KIIIα^25^. Although it is technically difficult to measure the local concentrations of PI4P in the PM of potentiated neurons, given the similar efficiency of PI4P and PI(4,5)P_2_ in promoting SNAP47 SNARE-mediated membrane fusion in *in vitro* reconstitution assays, it is plausible that at high concentrations, PI4P recruits SNAP47 to the PM and promotes assembly of the SNARE complex by enhancing the SNAP47–VAMP2 interaction, possibly via lipid binding-induced conformational changes. Conceivably, PI4P-enriched PM microdomains serves as hotspots for efficient fusion between cargo vesicles and the target membrane.

Of note, Ca^2+^ serves as a signaling molecule in all cell types. At presynaptic terminals, the Ca^2+^ sensor synaptotagmin on the surface of synaptic vesicles regulates rapid neurotransmitter release by triggering membrane fusion between docked vesicles and the PM ^59,60^. Ca^2+^ binding triggers electrostatic interactions of synaptotagmin with acidic phospholipids in the PM or the acidic surface of Q-SNAREs, or both ^61^. Moreover, PM PI(4,5)P_2_ enhances synaptotagmin activity to facilitate membrane fusion ^39^. In postsynaptic neurons, Syt-1 and Syt-7 serve as redundant Ca^2+^ sensors for Ca^2+^-induced exocytosis of AMPAR vesicles ^3^. As dendritic PM PI4P levels are upregulated by cytosolic Ca^2+^ in neurons undergoing synaptic potentiation ^25^, whether PI4P similarly modulates the function of synaptotagmins in exocytosis warrants further investigation.

## Materials and Methods

### Ethics statement

All animal experiments were approved by and performed in accordance with the guidelines of the Animal Care and Use Committee of Institute of Genetics and Developmental Biology, Chinese Academy of Sciences (Approval code: AP2022005). All animals were housed in standard mouse cages at 22–24°C on a 12 h light/dark cycle with access to food and water freely.

### Constructs and viruses

pCAG-SEP-GluA1 was a gift from Dr. Yong Zhang (Peking University, China) ^27^. The PH^PLCδ1^-EGFP construct was a gift from Tamas Balla (Addgene plasmid#51407; http://n2t.net/addgene:51407; RRID: Addgene_51407), the mCherry-PH^PLCδ1^ construct was a gift from Narasimhan Gautam (Addgene plasmid #36075; http://n2t.net/addgene:36075; RRID: Addgene_36075). pAKD-CMV-bGlobin-eGFP-H1-shRNA, pAKD-CMV-bGlobin-mCherry-H1-shRNA, and pAOV-CAMKIIα-mCherry-2A-3Flag were commercially obtained from OBiO Technology (Shanghai, China). The EGFP-PH^OSBP^ construct has been described previously ^62^. The mCherry-PH^OSBP^ construct was generated by replacing the EGFP coding sequence in the EGFP-PH^OSBP^ construct with mCherry. The expression constructs encoding dominant-negative forms of SNAP23 (lacking the C-terminal 8 residues) and other dominant-negative SNAP family members-including SNAP25, SNAP29, and SNAP47 (each lacking the C-terminal 20 residues) have been described previously ^30^. cDNA fragments encoding mouse SNAP47, Syntaxin-3 (Stx3), Syntaxin-4 (Stx4), and VAMP2 were amplified by RT-PCR from mouse brain total RNA and subsequently subcloned into pmCherry-C1, pmCherry-N1, or pEGFP-N1 as appropriate. Rat Stx3 and SNAP47 cDNA fragments were similarly amplified from rat brain total RNA by RT-PCR and cloned into pmCherry-C1. The pCMV-Flag-SNAP47 construct was generated by subcloning the full-length SNAP47 coding sequence, amplified from pCMV-SNAP47-mCherry, into the pCMV-Tag2B vector. The pCMV-Flag-SNAP47ΔPH construct was derived from pCMV-Flag-SNAP47 via deletion of amino acid residues 1–112. The pCMV-Flag-SNAP23 construct was generated by restoration of the C-terminal 8 residues into the dominant-negative SNAP23 backbone. For mammalian expression under the CAG promoter, pCAG-Flag-SNAP47, pCAG-Flag-SNAP47ΔPH, and pCAG-mCherry-SNAP47 were generated by subcloning the corresponding fragments from their pCMV-based counterparts into the pCAG vector. The pCAG-mCherry-SNAP47ΔPH construct was derived from pCAG-mCherry-SNAP47 by deletion of SNAP47 amino acid residues 1–112.

For bacterial expression, constructs for His-SNAP47, His-SNAP47ΔPH, His-Stx3, and His-SNAP23 were generated by subcloning the respective coding sequences amplified from pCMV-Flag-SNAP47, pCMV-Flag-SNAP47ΔPH, pCMV-Stx3-mCherry, and pCMV-Flag-SNAP23 into the pET-28a vector. His-SUMO-SNAP47 and His-SUMO-SNAP47ΔPH were generated by inserting the SUMO tag sequence into His-SNAP47 and His-SNAP47ΔPH, respectively. His-EEN1 has been described previously ^30^. GST-VAMP2 was generated by subcloning the VAMP2 coding sequence, amplified from pCMV-VAMP2-mCherry, into pGEX-4T-1.

To generate shRNA constructs, sense and antisense strands of each shRNA containing a 19-21 nt target sequence were annealed and inserted into pAKD-CMV-bGlobin-eGFP-H1-shRNA or pAKD-CMV-bGlobin-mCherry-H1-shRNA between the BglII and SalI sites. Target sequences are: for mouse SNAP47 (3′-UTR): 5′-ATAGCAATAGAATCAGCAGAGC-3′; mouse Stx3 #1 (3′-UTR): 5′-GCACCAATCAACTGTTTATTA-3′; mouse Stx3 #2: 5′-GGAAACACGGCTCAACATTGA-3′; mouse Stx4 #1: 5′-CCTGCGAGAGGAGATCAAA-3′; mouse Stx4 #2: 5′-AGACAATTCGGCAGACTAT-3′; rat Stx3: 5′-AGAGCATGGAGAAGCATATTG-3′; rat SNAP47: 5′-TCTCCTTATGAGGTCAGCATA-3′; for non-targeting control shRNA, which has no homology to known gene sequences: 5′-TCTCGCTTGGGCGAGAGTAAG-3′. The pAOV-CaMKIIα-mCherry-2A-SNAP47-3FLAG and pAOV-CaMKIIα-mCherry-2A-SNAP47ΔPH-3FLAG constructs were generated by subcloning the corresponding coding fragments amplified from pCAG-mCherry-SNAP47 and pCAG-mCherry-SNAP47ΔPH, respectively, into the pAOV-CaMKIIα-mCherry-2A-MCS backbone. All constructs generated in this study were verified by DNA sequencing.

Recombinant adeno-associated virus (rAAV) particles (packaging the following constructs: pAKD-CMV-bGlobin-eGFP-H1-shRNA-CTRL, pAKD-CMV-bGlobin-eGFP-H1-shRNA-SNAP47, pAOV-CaMKIIα-mCherry-2A-MCS-SNAP47-3FLAG, and pAOV-CaMKIIα-mCherry-2A-MCS-SNAP47ΔPH-3FLAG) were produced and titrated by OBiO Technology (Shanghai, China).

### Antibodies and reagents

Antibodies used in this study were mouse anti-GluA1 (MAB2263, Millipore, Billerica, MA, USA, 1:100 for immunofluorescence (IF)), rabbit anti-Flag (F7425, Sigma-Aldrich, St. Louis, MO, USA, 1:500 for IF, 1:3000 for WB), rabbit anti-GFP (598, MBL International, Woburn, MA, USA, 1:500 for IF, 1:3000 for WB), rabbit anti-RFP (PM005, MBL International, Woburn, MA, USA, 1:500 for IF, 1:3000 for WB), mouse anti-VAMP2 (sc-69706, Santa Cruz Biotechnology, Santa Cruz, CA, USA, 1:50 for IF), rabbit anti-GluA1 (13185, Cell Signaling Technology, Danvers, MA, USA, 1:200 for IF), rabbit anti-Myc (562, MBL International, Woburn, MA, USA, 1:3000 for WB), mouse anti-His (CW0285M, Cowin Biosciences, Jiangsu, China, 1:3000 for WB), mouse anti-β-Actin (A5441, Sigma-Aldrich, St. Louis, MO, USA, 1:3000 for WB), mouse anti-GAPDH (M171-3, MBL International, Woburn, MA, USA, 1:5000 for WB), rabbit anti-Rab5B (sc-598, Santa Cruz Biotechnology, Santa Cruz, CA, USA, 1:500 for WB), mouse anti-Na/K^+^ATPase (sc-48345, Santa Cruz Biotechnology, Santa Cruz, CA, USA, 1:500 for WB), rabbit anti-Calreticulin (MA532131, Thermo Fisher Scientific, Waltham, MA, USA, 1:2000 for WB). Alexa Fluor dye–conjugated secondary antibodies for IF staining were purchased from Invitrogen (Carlsbad, CA, USA), horseradish peroxidase (HRP)-conjugated secondary antibodies for WB were purchased from Bioss (Beijing, China).

Reagents used in this study were GSK-A1 (SYN-1219, SYNkinase, San Diego, CA, USA); Ovine cholesterol (700000P), Egg PC (840051P), Egg PE (840021P), Brain PS (840032P), Liver PI (840042P), 18:1 PI(3)P (850150P), 18:1 PI(4)P (850151P), 18:1 PI(5)P (850152P), 18:1 PI(3,4)P_2_ (850153P), 18:1 PI(3,5)P_2_ (850154P), 18:1 PI(4,5)P_2_ (850155P) and 18:1 PI(3,4,5)P_3_ (850156P) purchased from Avanti Research (Alabaster, Alabama, USA); PI diC8 (P-0008), PI(4)P diC8 (P-4008), PI(4,5)P_2_ diC8 (P-4508), PI(4)P diC16(P-4016), PI(4,5)P_2_ diC16(P-4516) purchased from Echelon biosciences (Salt Lake City, Utah, USA).

### Cell culture and transfection

HEK293T cells (Clontech #632180) were cultured in DMEM (C11995500BT, Gibco, Carlsbad, CA, USA) medium supplemented with 10 % (V/V) FBS (F0193, Sigma-Aldrich, St. Louis, MO, USA) at 37 °C with 5% CO_2_. Transient transfections of HEK293T cells with plasmid DNA were performed using Lipofectamine 2000 (11668019, Invitrogen, Carlsbad, CA, USA) according to the manufacturer’s protocol. Cells were harvested 24-48 h after transfection for analysis.

Primary hippocampal neurons were cultured as previously described ^25^. Briefly, hippocampi were dissected from newborn Sprague-Dawley rats or C57BL/6J mice, dissociated and incubated with 0.125% trypsin (SH30042.02, HyClone, Logan, UT, USA) at 37 °C for 18 min. Tissue was triturated in DMEM supplemented with 10% F-12 (319-085-CL, MultiCell Technologies, Woonsocket, RI, USA) and 10% FBS (11058021, Gibco, Carlsbad, CA, USA). Neurons were plated on poly-D-lysine (P6407, Sigma-Aldrich, St. Louis, MO, USA) coated coverslips in 24-well plates or 14 mm glass bottom dishes (D29-14-1.5-N, Cellvis, Mountain View, CA, USA) at a density of 2.5–3.0×10^4^ cells/well. The medium was replaced with the serum-free Neurobasal A (NB-A) (10888022, Gibco, Carlsbad, CA, USA) media supplemented with 2% B27 (17504044, Gibco, Carlsbad, CA, USA), 1% GlutaMAX (35050061, Gibco, Carlsbad, CA, USA) and 0.3% glucose 4 h after plating. Half of the media were replaced with fresh medium every 3 days until use. Primary hippocampal neurons were transfected using Lipofectamine LTX (15338100, Invitrogen, Carlsbad, CA, USA) according to the manufacturer’s instructions at 10-11 days in vitro (DIV) after plating and processed for experiments 6-8 days later.

### Chemical LTP induction (chemical LTP or cLTP)

Chemical induction of LTP with glycine was performed as previously described ^30^. Briefly, DIV 16-18 hippocampal neurons were treated with glycine (200 μM) in Mg^2+^-free extracellular solution (ECS) containing (in mM): 125 NaCl, 2.5 KCl, 2 CaCl_2_, 5 HEPES, 33 glucose, 0.2 glycine, 0.02 bicuculline, and 0.003 strychnine, pH 7.4 for 5 min to induce LTP and then incubated for 5 min or 30 min in without glycine in ECS. For live imaging, DIV 16-18 rat hippocampal neurons were treated with glycine (200 μM) in ECS containing (in mM): 125 NaCl, 2.5 KCl, 2 CaCl_2_, 25 HEPES, 10 glucose, 0.2 glycine, 0.0005 tetrodotoxin (TTX), 0.02 bicuculline, and 0.001 strychnine pH7.4 for 10 min to induce LTP.

### Immunofluorescence staining, confocal image acquisition and analysis

For surface GluA1 staining, hippocampal neurons at DIV 16-18 were fixed with 4% paraformaldehyde and 4% sucrose in PBS for 15 min at room temperature. After three PBS washes, cells were blocked with 5% normal goat serum (NGS) (ZLI-9022, ZSGB-BIO, Beijing, China) in PBS for 30 min and incubated overnight at 4°C with mouse anti-GluA1 antibody (1:100) in PBS containing 5% NGS. Following three PBS washes, cells were incubated with appropriate fluorescence-conjugated secondary antibodies for 1 h at room temperature. Neurons were then permeabilized with 1% BSA in PBS containing 0.4% Triton X-100 for 30 min at room temperature, followed by incubation with rat/rabbit anti-GFP (1:1500), anti-RFP (1:1500), or anti-Flag (1:1500) antibodies for 1 h. After three PBS washes, cells were incubated with appropriate fluorescence-conjugated secondary antibodies. For VAMP2-GluA1 colocalization analysis, after fixation, neurons were permeabilized and blocked with PBS containing1% BSA and 0.4% Triton X-100 for 30 min at room temperature, followed by labeling with rabbit anti-GluA1 (1:200) and mouse anti-VAMP2 (1:50) antibodies overnight at 4°C, then incubation with secondary antibodies conjugated with Alexa Fluor 488 and Alexa Fluor 647.

Confocal microscopy was performed on a Nikon EZ-A1 system (Nikon Corporation, Tokyo, Japan) equipped with a 100× Plan Apochromat VC oil immersion objective (numerical aperture 1.40). Image acquisition and subsequent analysis were conducted using NIS-Elements AR software (Nikon Corporation) or Fiji ImageJ (National Institutes of Health, USA).

### Protein expression and purification

His-SNAP47, His-SNAP47ΔPH, His-Stx3, His-EEN1, His-SUMO-SNAP47, His-SUMO-SNAP47ΔPH, His-SNAP23, and GST-VAMP2 were expressed in *Escherichia coli* BL21 (DE3) cells. Bacterial cultures were grown at 37°C in Luria-Bertani (LB) medium to an OD_600_ of 0.6, then induced with 0.4 mM IPTG and incubated for 20 h at 18°C. Cell pellets were harvested by centrifugation at 13,000 × g.

For purification of His-tagged proteins, cell pellets were resuspended in lysis buffer (50 mM NaH_2_PO_4_ pH 8.0, 300 mM NaCl, 10 mM imidazole, 1% Triton X-100, and 0.1 mM PMSF). Cells were lysed by sonication and clarified by centrifugation at 15,000 × g for 20 min at 4°C. The supernatant was incubated with Ni-NTA resin (70666-4, Millipore, Billerica, MA, USA) overnight at 4°C with rotation. The resin was washed three times with wash buffer (50 mM NaH_2_PO_4_, pH 8.0, 300 mM NaCl, 20 mM imidazole), and bound proteins were eluted three times with elution buffer (50 mM NaH_2_PO_4_, pH 8.0, 300 mM NaCl, 300 mM imidazole, 10% [v/v] glycerol).

For purification of GST-tagged proteins, cell pellets were resuspended in PBS supplemented with 0.2% Triton X-100 and 0.1 mM PMSF, then lysed by sonication. The clarified supernatant was incubated with glutathione Sepharose 4B beads (GE17-0756-01, Sigma-Aldrich, St. Louis, MO, USA) overnight at 4°C with rotation. The beads were subsequently washed three times with PBS containing 0.1% Triton X-100.

### Lipid blot overlay and liposome co-sedimentation assays

Lipid blot overlay assays were performed as previously described ^63^. Briefly, lipids were resuspended at 1 μg/μL in a chloroform/methanol/water solution (20:9:1, v/v/v) and spotted onto Hybond-C Extra nitrocellulose membranes (RPN303E, Amersham Bioscience, Little Chalfont, Buckinghamshire, UK). The membranes were blocked for 4 h at room temperature using TBS-T buffer (50 mM Tris-HCl pH 7.4, 150 mM NaCl, 0.05% Tween-20) supplemented with 3% fatty acid-free bovine serum albumin (BSA; BAH66, EquitechBio, Kerrville, TX, USA). After blocking, the membranes were incubated overnight at 4°C with recombinant His-tagged protein (1.5 μg/mL), then washed three times with TBS-T buffer. Subsequently, the blots underwent a second blocking step for 1 h using TBS-T containing 1% fatty acid-free BSA, followed by incubation with anti-His primary antibody and appropriate secondary antibody. Protein-lipid interactions were visualized using chemiluminescence detection (Minichemi 160, SAGECREATION, Beijing, China).

Liposome co-sedimentation assays were performed following established protocols ^62^. Liposomes were prepared with the following lipid composition: 50% phosphatidylcholine (PC), 25% phosphatidylethanolamine (PE), 20% phosphatidylserine (PS), and 5% phosphatidylinositol (PI), PI4P, or PI(4,5)P_2_ (molar ratio). The lipid mixtures were dried under vacuum in glass tubes for a minimum of 2 h. The resulting lipid films were resuspended in liposome buffer (150 mM NaCl, 20 mM HEPES pH 7.4, 1 mM DTT) and vortexed for 1 h at 37°C. This was followed by seven freeze-thaw cycles consisting of rapid freezing in liquid nitrogen and thawing in a 37°C water bath. The multilamellar vesicles were subsequently extruded 31 times through a 200 nm polycarbonate membrane using a mini-extruder (Avanti Polar Lipids) to generate uniform unilamellar liposomes. For the binding assay, liposomes (1 mM final concentration) were incubated with freshly purified recombinant proteins (2 μM) in 100 μL of liposome buffer for 20 minutes at 37°C. The samples were then subjected to ultracentrifugation at 140,000 × g for 30 minutes at 4°C. Both supernatant (unbound) and pellet (bound) fractions were collected and analyzed by SDS-PAGE and Coomassie blue staining. The binding ratios to PI and various phosphoinositides were quantified using ImageJ software (NIH).

### GST pull-down and co-immunoprecipitation (IP)

Co-immunoprecipitation (Co-IP) from HEK293T cells was performed as previously described ^63^. Briefly, HEK293T cells were lysed in ice-cold lysis buffer (50 mM Tris-HCl pH 8.0, 150 mM NaCl, 5 mM EDTA, 1% NP40) supplemented with protease inhibitors (1 mM PMSF, 10 μM leupeptin, 1 μg/mL pepstatin A, and 1 mM benzamidine) with gentle rotation for 30 min at 4°C. Cell lysates were clarified by centrifugation at 16,000 × g for 15 min at 4°C, and the supernatants were collected. For Flag- or GFP-tagged protein immunoprecipitation, the clarified lysates were incubated with anti-Flag (A2220, Sigma-Aldrich, St. Louis, MO, USA) or anti-GFP (SA070005, Smart-lifescience, Changzhou, Jiangsu, China) conjugated agarose beads overnight with gentle rotation at 4°C. The beads were subsequently washed three times with TBS-T buffer (50 mM Tris-HCl pH 7.4, 150 mM NaCl, 0.05% Tween-20). Bound proteins were eluted by boiling in 2× SDS-PAGE loading buffer at 100°C for 10 min and analyzed by SDS-PAGE and immunoblotting.

GST pull-down assays were performed as previously described ^64^. Briefly, 2 μg of His-tagged SNAP47 or SNAP47ΔPH and 2 μg His-tagged Stx3 were pre-incubated with 200 μM of phosphoinositides (PI diC8, PI(4)P diC8, or PI(4,5)P_2_ diC8) in binding buffer (10 mM HEPES pH 7.4, 150 mM NaCl, 1 mM EDTA, 0.5% NP-40, 0.5 mg/mL BSA, and 0.01% NaN_3_) for 3 h at room temperature to allow binary complex formation. The reaction mixture was then supplemented with 5 μg of GST-VAMP2 pre-immobilized on glutathione-Sepharose beads and incubated for an additional 3 h at room temperature to facilitate ternary complex assembly. Following incubation, the beads were washed three times with PBS containing 0.1% NP-40. Bound proteins were eluted by boiling in 2× SDS-PAGE loading buffer at 100°C for 10 min and analyzed by SDS-PAGE and immunoblotting.

### Ensemble liposome fusion assays

L-α-phosphatidylcholine (Egg PC, from egg, chicken), 1-palmitoyl-2-oleoyl-sn-glycero-3-phosphoethanolamine (POPE), 1,2-dioleoyl-sn-glycero-3-phospho-L-serine (DOPS), 1,2-dioleoyl-sn-glycero-3-phospho-(1’-myo-inositol-4’-phosphate) (ammonium salt) (PI(4)P), and L-α-phosphatidylinositol-4,5-bisphosphate (Brain, Porcine) (ammonium salt) (PI(4,5)P_2_) were obtained from Avanti Polar Lipids and dissolved in chloroform at a final concentration of 10 mg · ml^-1^ except for PI(4)P and PI(4,5)P_2_, which was dissolved in a mixture of chloroform:methanol 2:1 at a concentration of 1mg · ml^-1^. 1,1’-dioctadecyl-3,3,3’,3’-tetramethylindocarbocyanine (DiIC18(3)) (DiI) and 1,1’’-dioctadecyl-3,3,3’’,3’’-tetramethylindodicarbocyanine (DiDC18(5)) (DiD) were obtained from Molecular Probes and dissolved in ethanol at a concentration of 1 mg·ml^-1^. Lipids were mixed at the proper ratio as indicated below to a final concentration of 1 mM. Donor liposome (reconstituted with full-length VAMP2) contains 63.0% Egg PC, 20.0% POPE, 15.0% DOPS, and 2.0% DiI (molar ratio). Acceptor liposome (reconstituted with full-length Syntaxin 3) contains 58.0% Egg PC, 20.0% POPE, 15.0% DOPS, 5.0% PI(4)P/PI(4,5)P_2_, and 2.0% DiD (molar ratio). For no PI(4)P/PI(4,5)P_2_ supplement (no PIPs), the missing components were filled with Egg PC with corresponding molar ratio.

Lipid mixtures were dried under nitrogen flow and further incubated in vacuum for 16 h at room temperature (25°C) in photoprotective vacuum hood. Lipid films were resuspended in 100 μl TBS150 (20 mM Tris-Cl pH 8.0, 150 mM NaCl) supplied with 0.2 mM Tris (2-Tris (2-carboxyethyl) phosphine (TCEP, Sigma-Aldrich) and 1% (w/v) Sodium Cholate (Aladdin). Purified proteins were added to resuspended lipids with a protein-to-lipid ratio of 1:200. After incubation at 25°C for 1 h, lipid-protein mixtures were desalted using homemade Sephadex G-25 desalting column (8.3 ml bed volume). Prepared proteoliposomes were stored at 4°C in the dark before use. Liposome fusion assays were carried out using a FluoDia T70 fluorescence plate reader (PTI) equipped with 530/10 excitation filter, 580/10 and 667/10 emission filters at 37°C. Donor and acceptor liposomes were mixed at a concentration of 100 μM (total lipids) with addition of 5 μM recombinant SNAP47. Donor (DiI) and acceptor (DiD) fluorescence were monitored every 40 seconds. Liposome fusion signals were interpreted as the raw FRET efficiency (proximity ratio, *E*_PR_) between the donor (DiI) and acceptor (DiD):

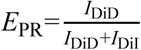

Where *I*_DiD_ and *I*_DiI_ are the fluorescence intensities of DiD and DiI under the excitation of 530/10 filter, respectively. All the experiments were independently repeated three times.

### TIRF live imaging acquisition and analysis

The TIRF-SIM system as described previously were utilized for live imaging ^65^. Rat hippocampal neurons were cultured on glass-bottom dishes (D29-10-1.5-N, Cellvis, Mountain View, CA, USA), transfected at DIV10-12 using Lipofectamine LTX (15338100, Invitrogen, Carlsbad, CA, USA), and imaged live at DIV16-18. To image GluA1 exocytosis, neurons were transfected with constructs expressing SEP-GluA1 and mCherry-PH^OSBP^ or mCherry-PH^PLCδ1^. To monitor GluA1 exocytosis and plasma membrane (PM) SNAP47 dynamics, neurons were co-transfected with SEP-GluA1 and mCherry-SNAP47-expressing plasmids and imaged before and after glycine application. To assess the role of the SNAP47 PH domain in PM targeting, neurons were transfected with mCherry-SNAP47 or mCherry-SNAP47ΔPH-expressing plasmid and similarly imaged upon glycine treatment. For detecting changes in PM phosphoinositide (PIP) and SNAP47 signals, neurons were co-transfected with a PIP biosensor and mCherry-SNAP47-expressing plasmids and imaged before and after glycine application. For drug treatment experiment, neurons were pre-treated with DMSO (vehicle control) or 10 nM GSK-A1 for 30 min prior to glycine-induced LTP, with the drug maintained throughout. To examine Stx3 dependence, neurons were co-transfected with mCherry-SNAP47 and either non-targeting (CTRL) shRNA or Stx3-targeting shRNA, then imaged to monitor SNAP47 dynamics upon glycine stimulation.

Images were acquired using a custom-built TIRF microscopy system equipped with an Olympus IX83 inverted microscope (100×/NA 1.49 oil, UAPON, Olympus, Japan) and an sCMOS camera (Dhyana 400BSI, Tucsen, China; Flash4.0 V2/Quest, Hamamastu, Japan). For dual-color imaging, 488 nm and 561 nm lasers (Genesis MX488-500 STM, Coherent; Jive 500-561 nm, Cobolt) were used for sequential excitation. An image splitter (Optosplit II, Cairn) was used before the sCMOS camera to split the emitted fluorescence into two channels. Images were acquired alternately at 488 nm and 561 nm with 50 ms exposure time using Tucsen Mosaic or HCImage software. Post-acquisition, red and green channel images were aligned using projective transformation. Prior to experiments, 100 nm fluorescent microspheres (blue/green/orange/dark red, T7279, Invitrogen) were imaged in both channels and their positions were mapped to calibrate channel alignment.

### Isolation of the plasma membrane by cell surface biotinylation and affinity capture with streptavidin-conjugated magnetic beads

Cell surface biotinylation was performed as previously described ^25^, with minor modifications adapted for each cell type. For primary neurons, cells were first washed twice with ice-cold Mg^2+^-free extracellular solution (ECS) containing (in mM): 125 NaCl, 2.5 KCl, 2 CaCl_2_, 5 HEPES, 33 glucose (pH 8.2) to remove residual culture medium and ensure full protonation of surface amine groups. Following removal of the wash buffer, cells were incubated with ice-cold Mg^2+^-free extracellular solution (pH 8.2) supplemented with 0.5 mg/mL EZ-Link Sulfo-NHS-LC-Biotin (Thermo Fisher Scientific, Waltham, MA, USA) for 30 min at 4°C on a rocking shaker. The biotinylation reaction was subsequently quenched by incubating cells with Mg^2+^-free extracellular solution containing 50 mM Tris-HCl (pH 8.2) for 10 min at 4°C on a rocking shaker. For HeLa cells, cells were washed twice with ice-cold phosphate-buffered saline (PBS, pH 7.4) prior to incubation with 0.5 mg/mL EZ-Link Sulfo-NHS-LC-Biotin dissolved in ice-cold PBS for 1 h at 4°C on a rocking shaker, followed by three washes with ice-cold PBS to remove unbound biotin reagent.

Cells were then lysed in ice-cold detergent-free lysis buffer (50 mM Tris-HCl, 10 mM HEPES pH 7.4, 150 mM NaCl, 0.5 mM EDTA, 0.25 M sucrose, supplemented with protease inhibitor cocktail) by passage through a 1 mL syringe needle at least 20 times to generate a uniform homogenate. The homogenate was subjected to two sequential low-speed centrifugation steps at 2,000 × *g* for 5 min at 4°C to remove cell debris and nuclei. The supernatant was subsequently centrifuged at 20,000 × *g* for 30 min at 4°C to pellet the membrane fraction. The resulting pellet was resuspended in lysis buffer and incubated with streptavidin-conjugated magnetic beads (SM017005, Smart Lifesciences, Changzhou, China) overnight at 4°C with gentle rotation. Beads were washed three times with lysis buffer lacking sucrose to remove non-specifically bound proteins. Biotinylated surface proteins were eluted by boiling in 2× SDS sample loading buffer and subsequently analyzed by SDS-PAGE and immunoblotting to determine the efficiency of biotinylation-affinity purification.

### Lipidomics analysis

Analyses of phosphoinositides (PIPs) and polar lipids were performed at LipidALL Technologies according to published protocols ^66,67^. Lipids were extracted from the plasma membranes isolated by surface biotinylation-streptavidin affinity purification using a modified version of the Bligh and Dyer’s method under acidic conditions to increase recovery of phosphoinositides ^68^. In brief, samples were incubated with an extraction solvent containing chloroform: methanol (1:1) + 0.5N HCl and 2 mM AlCl_3_ for 10 min at 1500 rpm. Internal standard cocktail containing PI3P-37:4, PI4P-37:4, PI5P-37:4, PI(3,4)P_2_-37:4, PI(3,5)P_2_-37:4, PI(4,5)P_2_-37:4 and PI(3,4,5)P_3_-37:4 from Avanti Polar Lipids was added into individual samples during extraction. At the end of incubation, deionized water was added to induce phase separation and the samples were centrifuged. The lower organic phase containing lipids were extracted and transferred to new tube. The extraction procedure was repeated for three rounds. The pooled lipid extract was divided between direct polar lipid analysis and derivatisation for analysis of PIPs, as elaborated below.

Direct polar lipid analysis was conducted on an ExionLC-AE coupled with Sciex QTRAP 7500 PLUS as reported previously ^69,70^. Separation of individual lipid classes of polar lipids by normal phase (NP)-HPLC was carried out using a TUP-HB silica column (i.d. 150×2.1 mm, 3 µm) with the following conditions: mobile phase A (chloroform:methanol:ammonium hydroxide, 89.5:10:0.5) and mobile phase B (chloroform:methanol:ammonium hydroxide:water, 55:39:0.5:5.5). MRM transitions were set up for analyses of various polar lipids. Individual lipid species were quantified by referencing to spiked internal standards. d_9_-PC32:0(16:0/16:0), d_9_-PC36:1p(18:0p/18:1), d_7_-PE33:1(15:0/18:1), d_9_-PE36:1p(18:0p/18:1), d_31_-PS(d_31_-16:0/18:1), d_7_-PA33:1(15:0/18:1), d_7_-PG33:1(15:0/18:1), d_7_-PI33:1(15:0/18:1), d_5_-CL72:8(18:2)4, d_7_-LPC18:1, d_7_-LPE18:1, C17-LPI, C17-LPA, C17-LPS, C17-LPG were obtained from Avanti Polar Lipids. GM3-d18:1/18:0-d_3_ was purchased from Matreya LLC. Free fatty acids were quantitated using d_31_-16:0 (Sigma-Aldrich) and d_8_-20:4 (Cayman Chemicals).

For PIPs analysis, lipid extract was derivatized with 2 M TMS-diazomethane in hexane according to a published protocol ^67^ and analyzed on a Shimadzu Nexera 30-AD HPLC coupled with Sciex TRIPLE QUAD 6500 PLUS. Isomers of PIPs from different classes were separated on a Daicel Chiralpak IB-U column (100 mm×3.0 mm, 1.6 μm) using 10 mM ammonium formate in water as mobile phase A, and methanol as mobile phase B ^71^. Endogenous PIPs were quantitated by referencing to the levels of internal standards added during lipid extraction.

### Stereotaxic injection of adeno-associated viruses into mouse brain

To suppress SNAP47 expression in defined brain regions, neonatal mice underwent stereotaxic injection of high-titer adeno-associated viruses (AAVs) carrying pAKD-CMV-bGlobin-eGFP-H1-SNAP47 shRNA (2.0 × 10¹² viral genomes/ml) into the hippocampal CA1 area within 24 h after birth. Neonates were anesthetized using hypothermia-induced analgesia by placement on ice for 4–5 min and secured in a custom-built ceramic stereotaxic apparatus, with lambda designated as the zero reference point for mediolateral and anteroposterior coordinates. Viral suspensions (10 nL per site) were delivered bilaterally into seven predefined locations per hemisphere using a microsyringe (Sutter Instrument, CA, USA) equipped with beveled glass micropipettes. Injection coordinates (in mm) were [X, Y, Z] = [1.2, 1.2, 1.4/1.0/0.6] and [1.5, 1.0, 1.7/1.3/0.9/0.5], targeting the hippocampal formation. For rescue experiments, age-matched littermates received co-injection of AAVs expressing SNAP47 shRNA and either AAV-CaMKIIα-mCherry-2A-MCS-SNAP47-3FLAG or AAV-CaMKIIα-mCherry-2A-MCS-SNAP47ΔPH-3FLAG (each at 2.0 × 10¹² viral genomes/ml). In these experiments, approximately 20 nL of viral mixture was administered at each injection site. Following surgery, pups were immediately returned to their home cages and allowed to recover under maternal care. Electrophysiological recordings were conducted approximately two weeks after viral delivery. Stereotaxic injection of viral particles into adult mouse brain was performed as previously described (Guo et al., 2022). Briefly, 8-week-old male mice (C57BL/6J) were anesthetized with tribromoethanol (330–390 mg/kg; T48402, Sigma-Aldrich, St. Louis, MO, USA) and placed in a stereotaxic apparatus. Following sterilization with iodophors and 75% (v/v) alcohol, a midline incision was made in the scalp between the ears, and bilateral cranial holes were drilled. Viral vectors were bilaterally injected into the hippocampal CA1 regions using a microinjection system (World Precision Instruments, Sarasota, FL, USA). The injection cocktail consisted of viral particles carrying pAKD-CMV-bGlobin-eGFP-H1-CTRL shRNA or pAKD-CMV-bGlobin-eGFP-H1-SNAP47 shRNA (0.6 μl, 2.0 × 10¹² viral genomes/ml) combined with those carrying pCaMKIIα-mCherry-2A-MCS-3FLAG, pCaMKIIα-mCherry-2A-MCS-SNAP47-3FLAG, or pCaMKIIα-mCherry-2A-MCS-SNAP47△PH-3FLAG (0.4 μl,

2.0 × 10¹² viral genomes/ml). Stereotaxic coordinates relative to bregma were as follows: anteroposterior (AP), −2.0 mm; mediolateral (ML), ±1.8 mm; dorsoventral (DV), −1.4 mm. The injection flow rate was maintained at 0.2 μl/min. The needle was maintained in place for 2 min before slow retraction. The scalp was then sutured, and the mice were kept on a heating pad during recovery.

### Electrophysiology

For preparation of acute hippocampal slices, mice were deeply anesthetized with tribromoethanol (330–390 mg/kg) and rapidly decapitated. Brains were quickly extracted and submerged in ice-cold, oxygenated (95% O_2_/5% CO_2_) sucrose-based cutting solution containing (in mM): 2.5 KCl, 25 NaHCO_3_, 1.25 NaH_2_PO_4_, 10 glucose, 210 sucrose, 1.3 sodium ascorbate, 0.5 CaCl_2_, and 7 MgSO_4_. Coronal hippocampal slices (300 μm thickness) were generated using a vibratome (Leica VT1200S; Leica Biosystems, Wetzlar, Germany). Slices were immediately transferred to standard artificial cerebrospinal fluid (ACSF) composed of (in mM): 119 NaCl, 2.5 KCl, 1.3 MgSO_4_, 2.5 CaCl_2_, 11 glucose, 26.2 NaHCO_3_, and 1 NaH_2_PO_4_ (pH 7.3–7.4), maintained at 33°C and continuously aerated with 95% O_2_/5% CO_2_. To isolate glutamatergic synaptic currents mediated by AMPA and NMDA receptors, 10 μM bicuculline (505875, Sigma-Aldrich, St. Louis, MO, USA) was included in the ACSF to block GABA_A_ receptors. After an incubation period of at least 30 min, slices were transferred to the recording chamber and maintained at room temperature (22–25°C). Whole-cell recordings were performed under continuous perfusion with oxygenated ACSF at room temperature. Neurons were visualized using infrared differential interference contrast optics on an upright microscope (BX51WI, Olympus, Tokyo, Japan) equipped with a 40× water-immersion objective. Fluorescently labeled CA1 pyramidal neurons were identified via epifluorescence illumination (pE-300, CoolLED Ltd, Andover, UK). Recordings were obtained from one AAV-infected neuron and a neighboring uninfected neuron within the same slice. Electrophysiological signals were acquired using a Multiclamp 700B amplifier (Molecular Devices, CA, USA), filtered at 2 kHz, and digitized at 10 kHz with a Digidata 1550B interface (Molecular Devices, CA, USA). All experiments were conducted in a blinded manner. The sample size indicated in each figure corresponds to the number of recorded neurons. Unless otherwise stated, reagents were obtained from Sigma-Aldrich. Data acquisition and analysis were performed using pClamp 10.6 software (Molecular Devices). Schaffer collateral–evoked AMPAR-mediated excitatory postsynaptic currents (AMPAR-eEPSCs), NMDAR-mediated excitatory postsynaptic currents (NMDAR-eEPSCs), and paired-pulse ratios (PPRs) were recorded using borosilicate glass electrodes (3–7 MΩ) filled with an internal solution containing (in mM): 135 CsMeSO_3_, 10 HEPES, 0.3 EGTA, 5 QX-314, 4 MgATP, 0.3 NaGTP, 8 NaCl, and 0.1 Spermine-Cl4 (pH 7.3).

Schaffer collaterals were stimulated using a bipolar electrode positioned in the stratum radiatum of the CA1 region. AMPAR-eEPSCs were recorded at a holding potential of −70 mV. NMDAR-eEPSCs were measured at +40 mV, 150 ms following stimulation, a time point at which AMPAR-mediated currents had fully decayed. Paired-pulse facilitation was assessed by delivering two stimuli separated by a 50-ms interval at −70 mV, and calculating the ratio of the second to the first peak response. Long-term potentiation (LTP) of AMPAR-mediated synaptic responses was induced by applying a theta-burst stimulation protocol at 2 Hz for 90 s while depolarizing the postsynaptic neuron to 0 mV. This protocol was initiated following a stable baseline recording of 3–5 min, and no later than 7 min after achieving whole-cell access to minimize intracellular washout. Synaptic responses were subsequently monitored for 45 min at −70 mV. All electrophysiological data were analyzed using two-tailed Student’s *t*-tests. Differences were considered statistically significant when *p* < 0.05. Data are presented as mean ± SEM.

### Animal behavioral tests

For animal behavior experiments, a 2-week interval was allowed between AAV viral injection of adult mouse brain and behavioral assessment to ensure optimal transgene expression and complete surgical recovery.

### Open field test

Mice were individually placed in an open-field apparatus and given 10 min of free exploration. Locomotor parameters (total distance traveled) and anxiety-related indices (percentage of distance and duration in the center zone) were automatically recorded and quantified (the Anilab System, AniLab Software and Instruments Co., Ltd, Ningbo, China).

### Rotarod

Prior to testing, mice underwent a training session in which they were placed on a rotarod apparatus rotating at a fixed speed of 4 rpm for 1 min of habituation. Mice that fell during this acclimation period were immediately repositioned on their original lane, and training continued until 1 min elapsed. Subsequently, during the test session, the rotarod was configured to accelerate uniformly from an initial velocity of 4 rpm to a final velocity of 40 rpm over a 5 min interval. The latency and the velocity to fall from the accelerating rod was automatically recorded for each mouse.

### Elevated plus maze

The elevated plus maze apparatus comprises four perpendicular arms positioned 50 cm above ground level: two arms are open (no walls) and two are enclosed (with vertical walls). Each mouse was placed on the central platform of the elevated plus maze facing an open arm.

Automated tracking software (the Anilab System, AniLab Software and Instruments Co., Ltd) recorded behavior for 5 min and calculated relevant indices. Between trials, mice were returned to home cages and the maze was cleaned with 70% ethanol.

### Y maze spontaneous alternation

The Y-maze is composed of three arms intersecting at 120° angles to form a Y-shaped configuration. Each mouse was gently positioned at the central intersection and allowed unrestricted exploration of all three arms during a 6 min test session. Alternations were defined as successive entries into three different arms on overlapping triplet sets (e.g., ABC, CAB, BCA). The spontaneous alternation percentage was calculated as: (number of alternations / [total arm entries - 2]) × 100.

### Contextual fear conditioning

Fear conditioning was performed in a standard conditioning apparatus (Harvard Apparatus, MA, USA). On the training day, each mouse was individually placed in the conditioning chamber and allowed to explore freely for 90 seconds. A 30-second auditory cue served as the conditioned stimulus (CS), immediately followed by a 0.9 mA foot shock lasting 2 seconds. After a 60-second inter-trial interval (ITI), a second 30-second auditory CS was delivered. Over the next four consecutive days, mice were re-exposed to the same conditioning chamber and allowed to explore freely for 90 seconds. A single 30-second CS was then presented, and freezing behavior during the CS was quantified as the percentage of time spent freezing. To prevent interference from residual olfactory cues, the chamber was cleaned with 70% ethanol between subjects. All freezing data were automatically acquired and analyzed using FREEZING software (Harvard Apparatus, MA, USA).

### Morris water maze

Morris water maze was performed in a 1.2 m diameter circular pool filled with opaque water (colored with non-toxic white paint) maintained at 19–23°C. Visual cues were placed at the four cardinal points on the inner pool walls. All movement trajectories were tracked using the Smart 3.0 system (Panlab SMART). The experiment included a hidden platform training phase and a subsequent probe test. For hidden platform training, a circular platform was submerged 1 cm below the surface in a fixed target quadrant. Mice received four daily trials, starting from different cardinal directions facing the wall. The time required to find the platform was recorded as the escape latency. Mice that failed to locate the platform within 60 s were guided to it and allowed to stay for 10 s to reinforce memory, in such cases, a latency of 60 s was recorded. The daily escape latency was calculated as the average of the four trials. In the probe test, the platform was removed. Mice were introduced from the point furthest from the target quadrant, and the number of crossings over the original platform location was recorded during a 60 s session.

### Statistics and reproducibility

Unless otherwise stated, all data are presented as mean ± standard error of the mean (SEM) from at least three independent biological replicates. Statistical analyses were performed using GraphPad Prism 5 (GraphPad Software, San Diego, CA, USA). For comparisons between two groups, two-tailed unpaired Student’s t-test was used unless otherwise stated. For comparisons among three or more groups, one-way analysis of variance (ANOVA) followed by Dunnett’s or Tukey’s post hoc multiple comparison test was applied to determine statistical significance. Differences were considered statistically significant at *p* < 0.05.

## Resource availability

### Lead contact

Requests should be directed to Jia-Jia Liu.

### Materials availability

All unique/stable reagents generated in this study are available from the lead contact with a complete materials transfer agreement.

### Data and code availability

All relevant data and details of resources can be found within the article and its supplementary information.

This paper does not report any original code.

## Acknowledgements

We thank Dr. Pingyong Xu (Institute of Biophysics, Chinese Academy of Sciences) for expression constructs and access to TIRF microscopy. This work was supported by funding from the National Natural Science Foundation of China (32270730, 32570819, 31530039, 31921002 to J-J. Liu, 32271005 to Y. Yang, 32225024 to C. Ma, and 32401035 to S. Wang), National Key R&D Program of China, Ministry of Science and Technology of China (2025YFA1804600 to J-J. Liu), the Natural Science Foundation of Beijing Municipality (1S24087 to J-J. Liu), and the Science and Technology Projects of Xizang Autonomous Region (XZ202501ZY0120 to L. Chen).

## Author contributions

Conceptualization, J-J. Liu, C. Ma, Y. Niu, and L. Chen; Investigation, J. Xi, S. Wang, D. Pan, Y. Yang, R. Mao and W. He; Formal analysis, J. Xi, S. Wang, D. Pan, and S.M. Lam; Visualization, J. Xi, S. Wang, D. Pan, Y. Yang, and J-J. Liu; Writing, J. Xi, S. Wang, D. Pan and J-J. Liu; Funding acquisition, J-J. Liu, Y. Yang, C. Ma, S. Wang and L. Chen; Supervision, J-J. Liu, C. Ma and G. Shui; Project administration, J-J. Liu.

## Declaration of Interests

The authors declare no competing interests.

## Supplementary figures and captions for supplementary videos

**Fig. S1.**
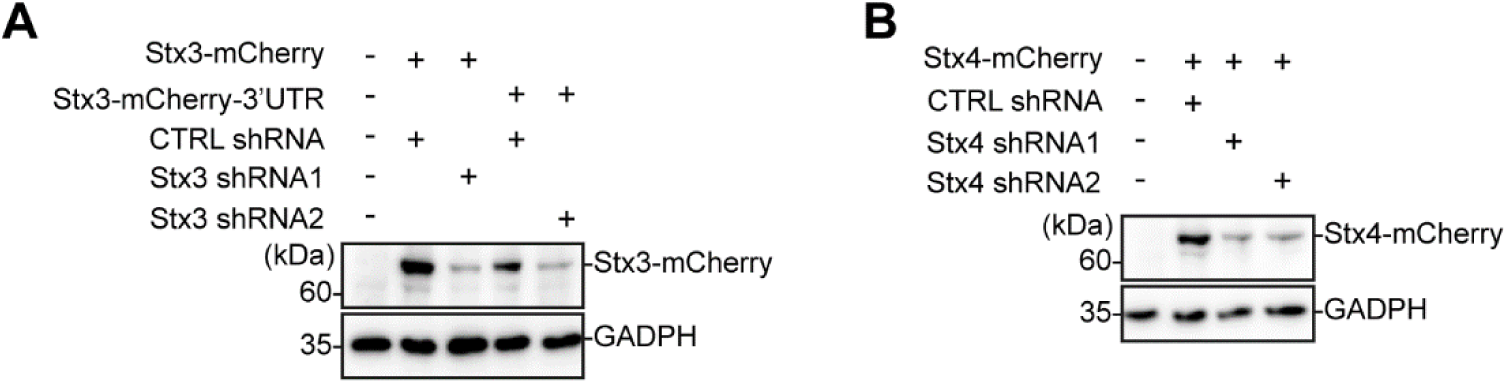
Knockdown efficiency of shRNAs targeting mouse Stx3 and Stx4. (A) HEK293T cells were co-transfected with constructs expressing non-targeting control shRNA (CTRL shRNA) or shRNA targeting mouse Stx3 (Stx3 shRNA1 targeting the coding sequence or Stx3 shRNA2 targeting the 3′-UTR of Stx3) and Stx3-mCherry or Stx3-mCherry-3′UTR. Cells were lysed and analyzed for Stx3-mCherry expression by western blotting with anti-RFP antibody. GAPDH served as loading control. (B) Same as (A) except that cells were co-transfected with constructs expressing either a non-targeting control shRNA (CTRL shRNA) or shRNA targeting mouse Stx4 (Stx4 shRNA1 or Stx4 shRNA2) and Stx4-mCherry.

**Fig. S2.**
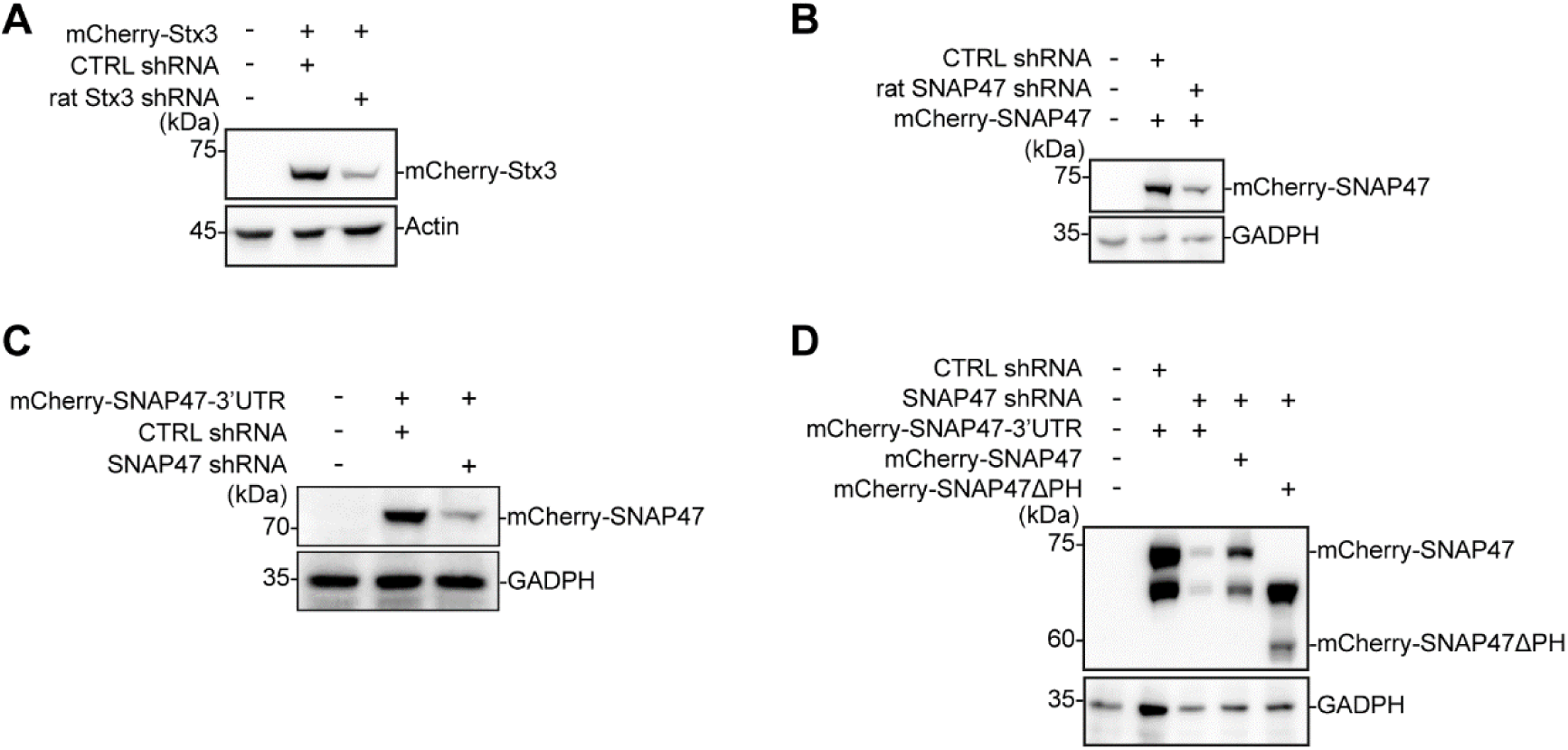
Knockdown efficiency of rat Stx3- and murine SNAP47-targeting shRNAs, and expression of RNAi-resistant SNAP47. (A) HEK293T cells were co-transfected with constructs expressing either a non-targeting control shRNA (CTRL shRNA) or shRNA targeting rat Stx3 (rat Stx3 shRNA) and mCherry-Stx3. Cell lysates were analyzed for mCherry-Stx3 expression by western blotting with anti-RFP antibody. Actin served as loading control. (B) Same as (A) except that cells were co-transfected with constructs expressing either a non-targeting control shRNA (CTRL shRNA) or shRNA targeting rat SNAP47 (rat SNAP47 shRNA) and mCherry-SNAP47. GAPDH served as loading control. (C) Same as (A) except that cells were co-transfected with constructs expressing either a non-targeting control shRNA (CTRL shRNA) or shRNA targeting the 3’UTR of mouse SNAP47 (SNAP47 shRNA) and mCherry-SNAP47-3’UTR. GAPDH served as loading control. (D) Same as (A) except that cells were co-transfected with constructs expressing non-targeting control shRNA (CTRL shRNA) and mCherry-SNAP47-3′UTR, or shRNA targeting the 3’UTR of mouse SNAP47 (SNAP47 shRNA) and mCherry-SNAP47-3′UTR, mCherry-SNAP47, or mCherry-SNAP47ΔPH. GAPDH served as loading control.

**Fig. S3.**
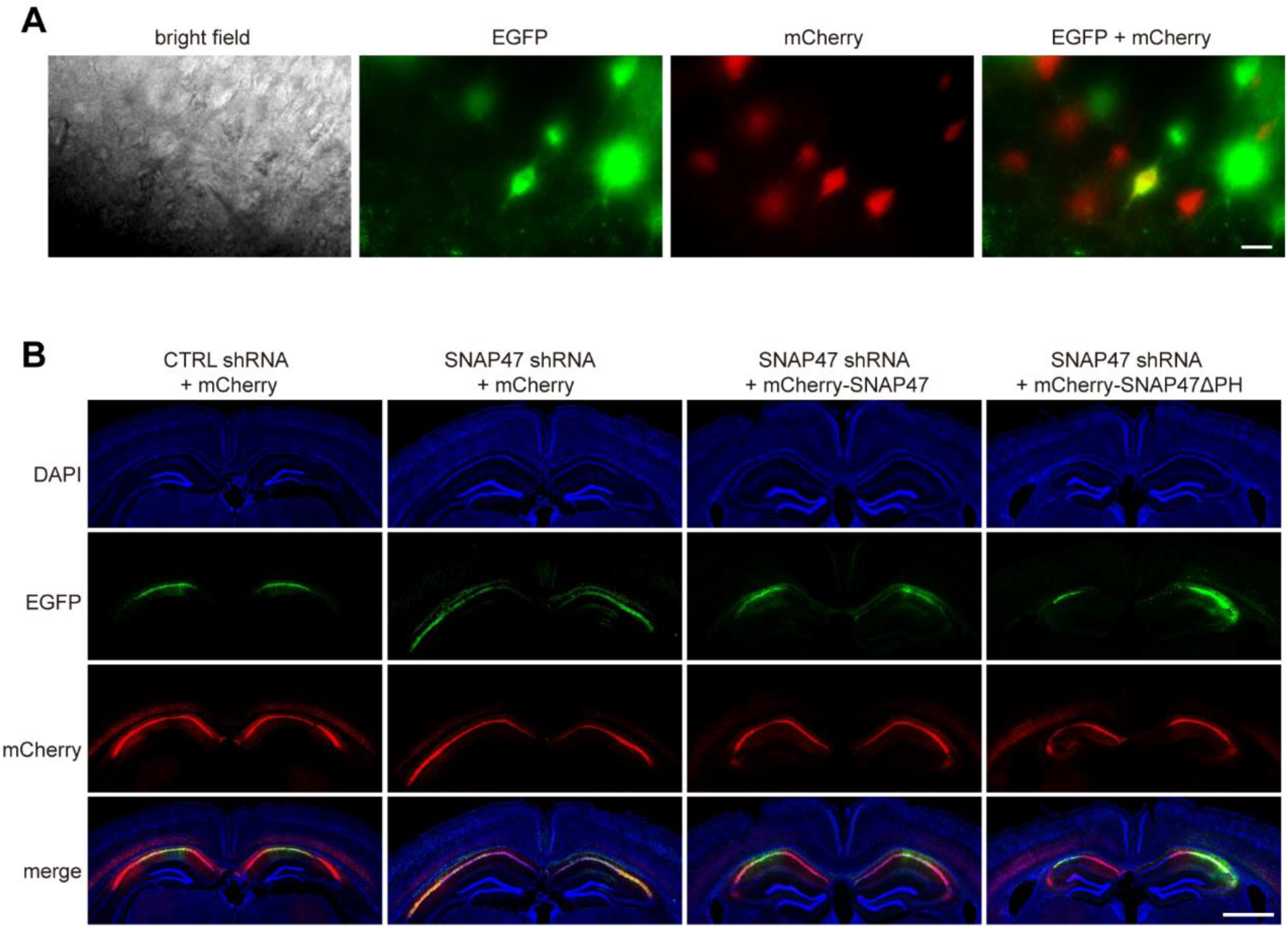
AAV expression in mouse brain. (A) AAVs co-expressing EGFP and SNAP47 shRNA and AAVs expressing mCherry, mCherry-SNAP47, or mCherry-SNAP47ΔPH were stereotaxically co-injected into the hippocampal CA1 region of P0 mice. Acute hippocampal slices were prepared between P14 and P21 for electrophysiological analyses. Representative images illustrate AAV-infected neurons in acute slices used for whole-cell patch-clamp recordings. Neurons co-expressing EGFP and mCherry (appearing yellow) were selected for analysis. Scale bar, 40 μm. (B) Eight-week-old male mice were stereotaxically injected with AAVs (CTRL shRNA + mCherry, SNAP47 shRNA + mCherry, SNAP47 shRNA + mCherry-SNAP47, or SNAP47 shRNA + mCherry-SNAP47ΔPH) in the hippocampal CA1 region. Shown are representative coronal brain sections depicting bilateral injections and expression of AAV. Scale bar, 1 mm.

**Figure S4.**
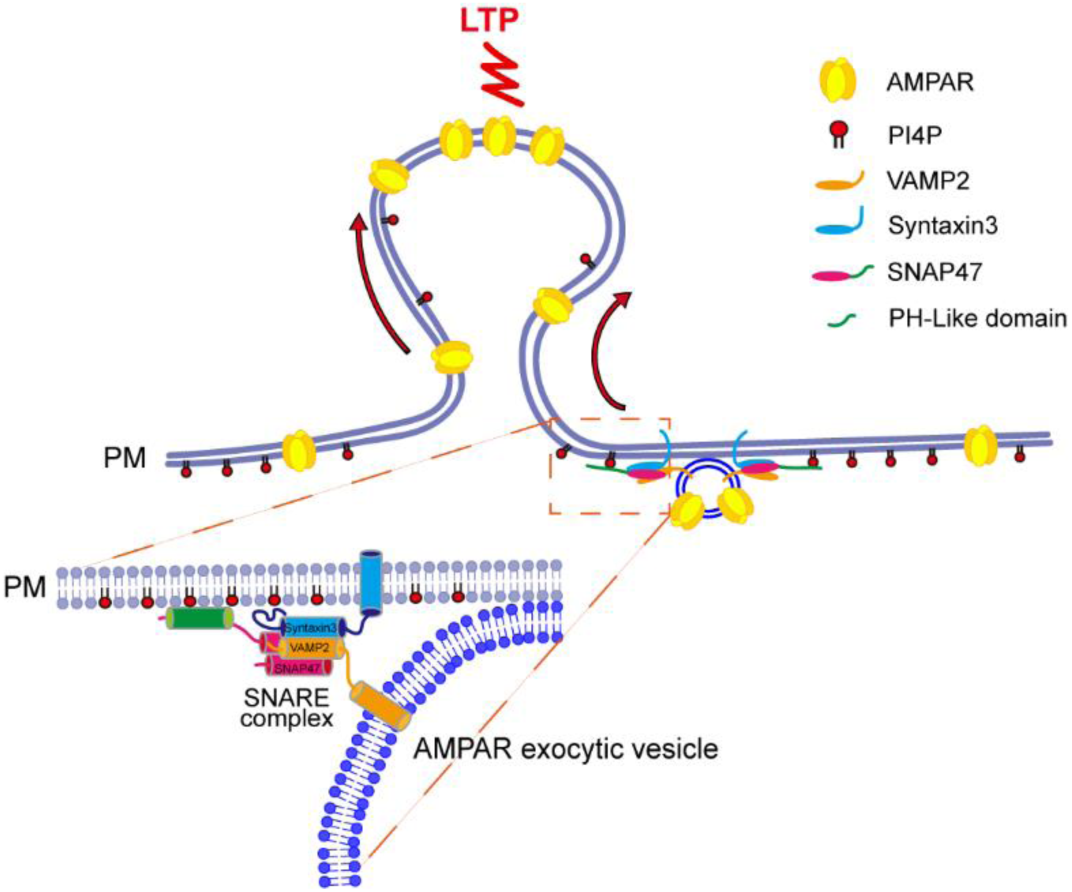
Model for PI4P-regulated AMPAR exocytosis. The SNAP47–PI4P interaction mediates activity-induced AMPAR exocytosis. During NMDAR-mediated LTP, PI4P enriched in the dendritic PM recruits SNAP47 and promotes the assembly of the SNAP47‒Syntaxin3‒VAMP2 SNARE complex, facilitating the fusion between AMPAR transport vesicles and the PM.

**Table S1. Molecular Fractions of individual PIP species to total PIs**

## Captions for supplementary videos

**Video 1. Activity-induced exocytosis of SEP-GluA1 at PI4P-enriched PM microdomains.** Cultured rat hippocampal neurons were co-transfected with constructs expressing SEP-GluA1 and either the PI4P probe mCherry-PH^OSBP^ or the PI(4,5)P_2_ probe mCherry-PH^PLCδ1^, and imaged live by TIRF microscopy before and after glycine application. Images were acquired at 5 frame/sec. Video plays at 200 frames/s. Scale bar, 2 μm.

**Video 2. Activity-induced PM recruitment of SNAP47 requires the PH-lie domain.** Cultured rat hippocampal neurons were transfected with constructs expressing either mCherry-SNAP47 or mCherry-SNAP47ΔPH and imaged live by TIRF microscopy before and after glycine application. Images were acquired at 0.5 frame/sec. Video plays at 30 frames/sec. Scale bar, 3 μm.

**Video 3. Activity-induced PM recruitment of SNAP47 requires PI4P.** Cultured rat hippocampal neurons were transfected with construct expressing mCherry-SNAP47 and imaged live by TIRF microscopy before and after glycine application. Cells were treated with DMSO (vehicle control) or the PI4KIIIα inhibitor GSK-A1. Images were acquired at 0.5 frame/sec. Video plays at 30 frames/s. Scale bar, 3 μm.

**Video 4. Neuronal activity induces recruitment of SNAP47 to PI4P-enriched PM microdomains.** Cultured rat hippocampal neurons were co-transfected with constructs expressing mCherry-SNAP47 and either the PI4P probe EGFP-PH^OSBP^ or the PI(4,5)P_2_ probe PH^PLCδ1^-EGFP, and imaged live by TIRF microscopy before and after glycine application. Images were acquired at 0.5 frame/sec. Video plays at 30 frames/s. Scale bar, 2 μm.

**Video 5. Silencing Stx3 expression does not inhibit activity-induced PM recruitment of SNAP47.** Cultured rat hippocampal neurons were co-transfected with constructs expressing mCherry-SNAP47 and non-targeting control shRNA (CTRL shRNA) or Stx3-targeting shRNA (Stx3 shRNA), and imaged live by TIRF microscopy before and after glycine application. Images were acquired at 0.5 frame/sec. Video plays at 30 frames/sec. Scale bar, 3 μm.

**Video 6. Activity-induced SEP-GluA1 exocytosis at SNAP47-localized PM microdomains.** Cultured rat hippocampal neurons were co-transfected with constructs expressing SEP-GluA1 and mCherry-SNAP47, and imaged live by TIRF microscopy before and after glycine application. Images were acquired at 0.5 frame/sec. Video plays at 30 frames/sec. Scale bar, 2 μm.

**Video 7. Silencing SNAP47 expression inhibits activity-induced SEP-GluA1 exocytosis.** Cultured rat hippocampal neurons were co-transfected with constructs expressing SEP-GluA1 and non-targeting control shRNA (CTRL shRNA) or SNAP47-targeting shRNA (SNAP47 shRNA), and imaged live by TIRF microscopy before and after glycine application. Images were acquired at 1 frame/sec. Video plays at 30 frames/sec. Scale bar, 3 μm.

## Notes

### Competing Interest Statement

The authors have declared no competing interest.

